# Molecular basis of interactions between CaMKII and α-actinin-2 that underlie dendritic spine enlargement

**DOI:** 10.1101/2022.12.04.519035

**Authors:** Ashton J. Curtis, Jian Zhu, Christopher J. Penny, Matthew G. Gold

**Affiliations:** Department of Neuroscience, Pharmacology & Physiology, University College London, Gower Street, LONDON, WC1E 6BT, UK

## Abstract

Ca^2+^/calmodulin-dependent protein kinase II (CaMKII) is essential for long-term potentiation (LTP) of excitatory synapses that is linked to learning and memory. In this study, we focused on understanding how interactions between CaMKIIα and the actin crosslinking protein α-actinin-2 underlie long-lasting changes in dendritic spine architecture. We found that association of the two proteins was unexpectedly elevated following stimulation of NMDA receptors to trigger structural LTP in primary hippocampal neurons. Furthermore, disruption of interactions between the two proteins prevented the accumulation of enlarged mushroom-type dendritic spines following NMDA receptor activation. α-actinin-2 binds to the regulatory segment of CaMKII. Calorimetry experiments, and a crystal structure of α-actinin-2 EF hands 3 and 4 in complex with the CaMKII regulatory segment, indicate that the regulatory segment of autoinhibited CaMKII is not fully accessible to α-actinin-2. Pull-down experiments show that occupation of the CaMKII substrate binding groove by GluN2B markedly increases α-actinin-2 access to the CaMKII regulatory segment. Overall, our study provides new mechanistic insight into the molecular basis of structural LTP and reveals an added layer of sophistication to the function of CaMKII.

## Introduction

Changes in synaptic connections between neurons are fundamental to learning and memory (Citri & Malenka, 2008). Calmodulin-dependent protein kinase II (CaMKII) plays a central role in long-term potentiation (LTP) of excitatory synapses following influxes of Ca^2+^ into postsynaptic spines (Hell, 2014). Activation of CaMKII by Ca^2+^/CaM leads to phosphorylation of postsynaptic proteins including AMPA-type glutamate receptors which has the overall effect of increasing postsynaptic responsiveness to glutamate release (Anggono & Huganir, 2012). CaMKII also serves a structural function in LTP (Hojjati et al., 2007) by nucleating networks of protein-protein interactions through its modular oligomeric structure (Bhattacharyya et al., 2020), consistent with its extremely high abundance in dendritic spines (Erondu & Kennedy, 1985). CaMKII holoenzymes form through oligomerization of the C-terminal hub domains of twelve subunits (Chao et al., 2011) (Myers et al., 2017). The hub domains assemble into a two-tiered central ring from which the N-terminal kinase domains radiate (Chao et al., 2011; Myers et al., 2017). Binding of Ca^2+^/CaM to a central regulatory segment releases the segment from the kinase domain enabling access to substrates and interaction partners (Yasuda et al., 2022). The CaMKII kinase domain has many documented substrates and binding partners (Özden et al., 2020). Formation of a highly-stable complex between the CaMKII kinase domain and the C-terminal tail of NMDA receptor (NMDAR) GluN2B subunits is thought to be critically important for learning and memory (Sanhueza et al., 2011). Unlike the promiscuous kinase domain, the regulatory segment of CaMKIIα interacts only with α-actinins (Dhavan et al., 2002; Walikonis et al., 2001) besides CaM, while the sole binding partner of the hub domain is densin-180 (Strack, Robison, et al., 2000). Understanding how and when the regulatory and hub regions of CaMKII engage in these interactions is essential for a complete understanding of the structural role of CaMKII in LTP (Hell, 2014).

α-actinins are structural proteins that form antiparallel rod-like dimers (Almeida Ribeiro Jr. et al., 2014). In the heart, they are a key component of Z-discs where they crosslink actin filaments to titin (Young et al., 1998). In dendritic spines, α-actinins serve a more complex function with many additional binding partners including CaMKII, NMDARs (Wyszynski et al., 1997), PSD-95 (Matt et al., 2018), CaV1.2s (Hall et al., 2013), and densin-180 (Walikonis et al., 2001). Three of the four α-actinin isoforms have been detected in the postsynaptic density (Walikonis et al., 2000). The most abundant is α-actinin-2, which localizes to dendritic spines of excitatory synapses (Rao et al., 1998; Wyszynski et al., 1998) where it interacts with the CaMKII regulatory segment independent of Ca^2+^ through its third and fourth EF hands close to its C-terminus (gold, **Fig. 1a**) (Jalan-Sakrikar et al., 2012). Association with actinin only weakly stimulates the activity of CaMKII towards certain substrates (Jalan-Sakrikar et al., 2012) therefore the interaction is thought to serve a structural role. Immunogold labelling shows that α-actinin-2 is present within the postsynaptic density (Wyszynski et al., 1998). Knockdown or overexpression of α-actinin-2 in cultured hippocampal neurons leads to defects in spine formation, with α-actinin-2 knockdown breaking the link between NMDAR activation and spine enlargement (Hodges et al., 2014) – a process known as structural LTP.

**Figure 1.**
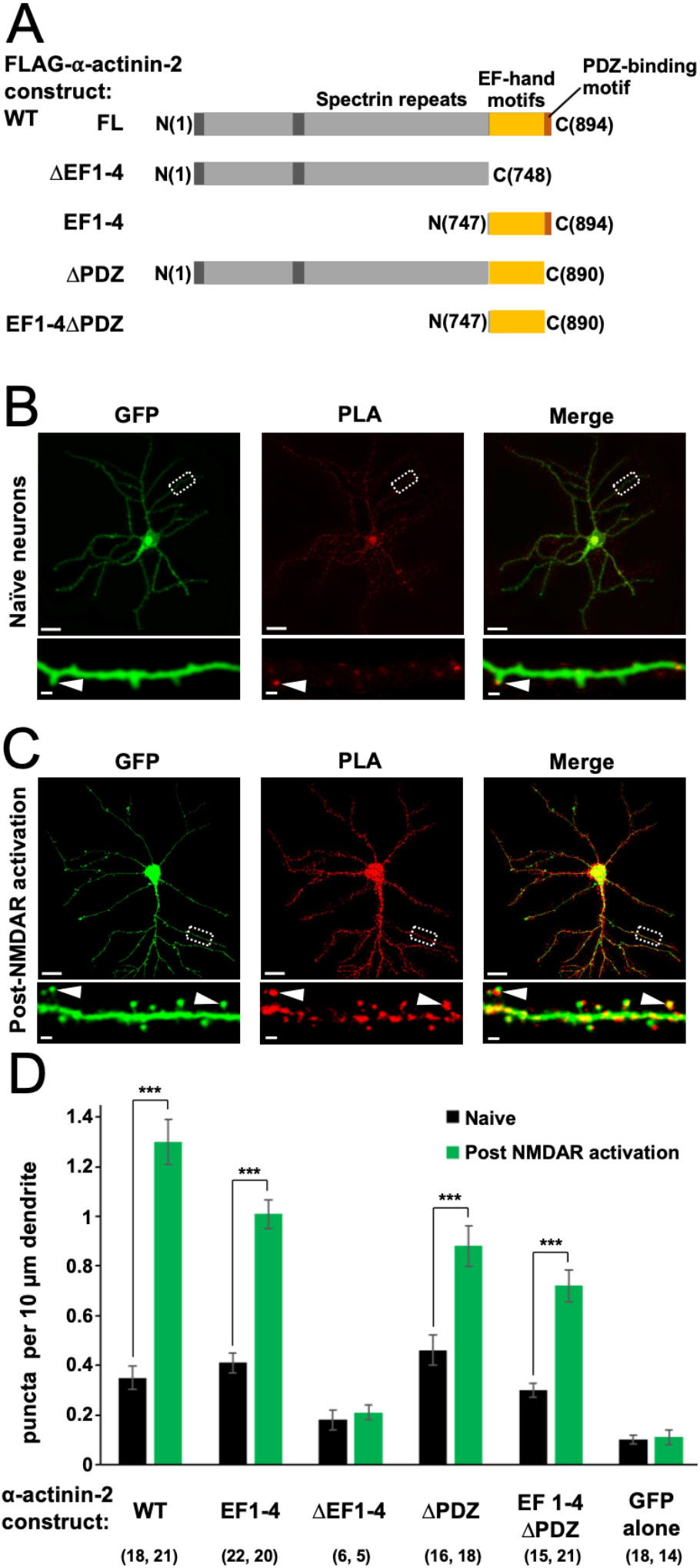
Changes in association of α-actinin-2 and CaMKII following NMDAR activation. (A) Topologies of α-actinin-2 constructs expressed in neurons. (B) and (C) show anti-GFP immunofluorescence (left column) and anti-CaMKIIα/anti-FLAG PLA puncta (middle column) in primary hippocampal neurons expressing FLAG-α-actinin-2 WT with GFP either before (B) or after (C) NMDAR activation. Scale bars are 20 *μm* (whole neuron images) and 1 *μm* (dendrite close-ups). (D) Quantitation of PLA puncta per 10 *μm* dendrite before (black) and after (green) NMDAR activation for the full range of α-actinin-2 constructs. Data are presented as the mean ± standard error (SE), and were collected from at least three animals with the exception of ΔEF1-4 (one animal). The number of neurons analysed for each construct is shown in parentheses.

Given that α-actinin-2 is known to bind both CaMKII and actin filaments – and is required for spine enlargement – it would be logical if it bridged the two to support the structure of enlarged spine heads that accumulate following LTP. However, this model is at odds with the observation that Ca^2+^/CaM (which activates CaMKII during LTP induction) competes with α-actinin-2 for binding to the CaMKII regulatory segment *in vitro* (Jalan-Sakrikar et al., 2012). Furthermore, previous modelling studies suggest that the regulatory segment is fully accessible to α-actinin-2 in inactive CaMKII (Jalan-Sakrikar et al., 2012), undermining the notion that the two associate following CaMKII activation to bring about structural LTP. In this study, we have focused on reconciling these apparently contradictory observations. We employed *in situ* labelling in primary hippocampal neurons to reveal changes in association of CaMKIIα and α-actinin-2 following a NMDAR stimulation protocol that triggers structural LTP. We also examined the effect of disrupting the actinin-CaMKII interface on spine head enlargement triggered by NMDAR activation. We used isothermal titration calorimetry and crystallography to determine whether the CaMKII kinase domain occludes access of α-actinin-2 to the regulatory segment in the inactive enzyme, and we have identified a potential mechanism for increasing access to the regulatory segment following induction of LTP. This combination of experiments has enabled us to put forward a more coherent molecular model of structural LTP.

## Results

### Association of CaMKII and α-actinin-2 is elevated following spine head enlargement

To investigate actinin association with CaMKII in neurons, we utilized proximity ligation assays (PLAs). We developed a set of vectors for mammalian expression of different fragments of α-actinin-2 (***Figure 1***A) in tandem with GFP. Each α-actinin-2 construct bears an N-terminal FLAG tag so association with CaMKII can be monitored by PLA using paired anti-FLAG and anti-CaMKIIα antibodies. We transfected each construct in rat primary hippocampal neurons after 10 days *in vitro* (DIV10), and fixed the mature neurons on DIV14 for imaging anti-GFP immunofluorescence and α-actinin-2-CaMKIIα PLA puncta. For wild-type (WT) α-actinin-2, PLA puncta were visible in dendritic spines in unstimulated neurons (***Figure 1***B, arrow) at a density of 0.35±0.05 puncta per 10 μm dendrite (**Figure *1***D) – 3.6-fold higher (*p*=2.2×10^-5^) than in unstimulated neurons expressing GFP alone (***Figure 1-figure supplement 1***A). To investigate the notion that interactions between actinin and CaMKII stabilize the structure of potentiated spines, we employed a chemical model of LTP that uses glycine to activate NMDARs to triggers spine head enlargement (Fortin et al., 2010; Shahi & Baudry, 1993). This is considered a relatively realistic chemical model for LTP (Fortin et al., 2010) that mimics other features including an increase in mini excitatory postsynaptic current amplitude (Lu et al., 2001) following stimulation. Anti-GFP immunofluorescence indicated that many mushroom-type spines had formed 4 hours after NMDAR activation, as expected (***Figure 1***C, left panel). This was associated with a 3.7-fold increase (*p*=1.6×10^-10^) in puncta in neurons expressing WT α-actinin-2 (***Figure 1***C&D). Bright puncta were localised to enlarged mushroom-type spines (***Figure 1***C, arrows), consistent with a role for α-actinin-2-CaMKIIα interactions in stabilizing the structure of potentiated spines.

Next, we compared PLA puncta formation in neurons expressing different fragments of α-actinin-2 (***Figure 1***A). Consistent with a key role for α-actinin-2 EF hands in mediating interactions with CaMKIIα (Jalan-Sakrikar et al., 2012; Robison, Bass, et al., 2005), expression of a construct limited to the EF hand motifs (‘EF1-4’) yielded similar results to those obtained with full-length α-actinin-2 with a puncta density of 0.41±0.04 per 10 μm dendrite rising to 1.01±0.06 (*p*=6.8×10^-11^) following NMDAR activation (***Figure 1**D* & ***Figure 1-figure supplement 1***B). Deletion of the EF hands (‘ΔEF1-4’) brought puncta density close to baseline levels irrespective of NMDAR activation with 0.18±0.04 and 0.21±0.03 puncta/10 μm dendrite before and after NDMAR activation, respectively (***Figure 1***D & ***Figure 1-figure supplement 1***C). The last four amino acids of α-actinin-2 (ESDL) are capable of binding to the PDZ domain of densin-180 (Walikonis et al., 2001) *in vitro*, and cooperative interactions between the three proteins in ternary complexes have been put forward as potentially important for supporting the structure of dendritic spines (Robison, Bass, et al., 2005; Walikonis et al., 2001). Expression of FLAG-α-actinin-2 lacking the last four amino acids (‘ΔPDZ’) generated 0.46±0.06 PLA puncta per 10 μm in naïve synapses, rising to 0.88±0.08/10 μm (*p*=2.9×10^-4^) following NMDAR activation (***Figure 1**D* & ***Figure 1-figure supplement 1***D). Similar responses were obtained with an EF hand construct lacking the PDZ motif (‘EF1-4ΔPDZ’) with 0.30±0.03 puncta per 10 μm dendrite rising to 0.72±0.07/10 μm (*p*=1.1×10^-5^) following NMDAR activation (***Figure 1**D* & ***Figure 1-figure supplement 1***E). Overall, we saw a mild attenuation of puncta formation when the PDZ motif was absent, most apparent in the context of the full-length protein following NMDAR activation (1.48-fold reduction in puncta density, *p*=0.0017). Nevertheless, our data indicate that the actinin EF hands are sufficient for robust α-actinin-2 – CaMKII interactions, and this region underlies the marked elevation in interaction of the two proteins in enlarged spines.

### Disruption of CaMKII-actinin interaction prevents spine enlargement following NMDAR activation

We noticed during PLA experiments that dendrites of neurons expressing α-actinin-2 EF1-4 seemed to exhibit fewer mushroom type spines following NMDAR activation (***Figure 1-figure supplement 1***B) than those expressing either GFP alone (***Figure 1-figure supplement 1***A) or the full-length α-actinin-2 construct (***Figure 1***C). The α-actinin-2 EF1-4 construct (**Figure 1***A*) lacks elements including the actin-binding domain and therefore can be expected to operate as a disruptor of actinin-mediated linking of CaMKII to the actin cytoskeleton. To investigate this disruptor effect further, we compared numbers of stubby (red), thin (amber) and mature mushroom-type (green) spines in neurons expressing either GFP alone (***Figure 2***A), GFP with α-actinin-2 EF1-4 WT (***Figure 2***B), or GFP with α-actinin-2 EF1-4 L854R (***Figure 2***C).

**Figure 2.**
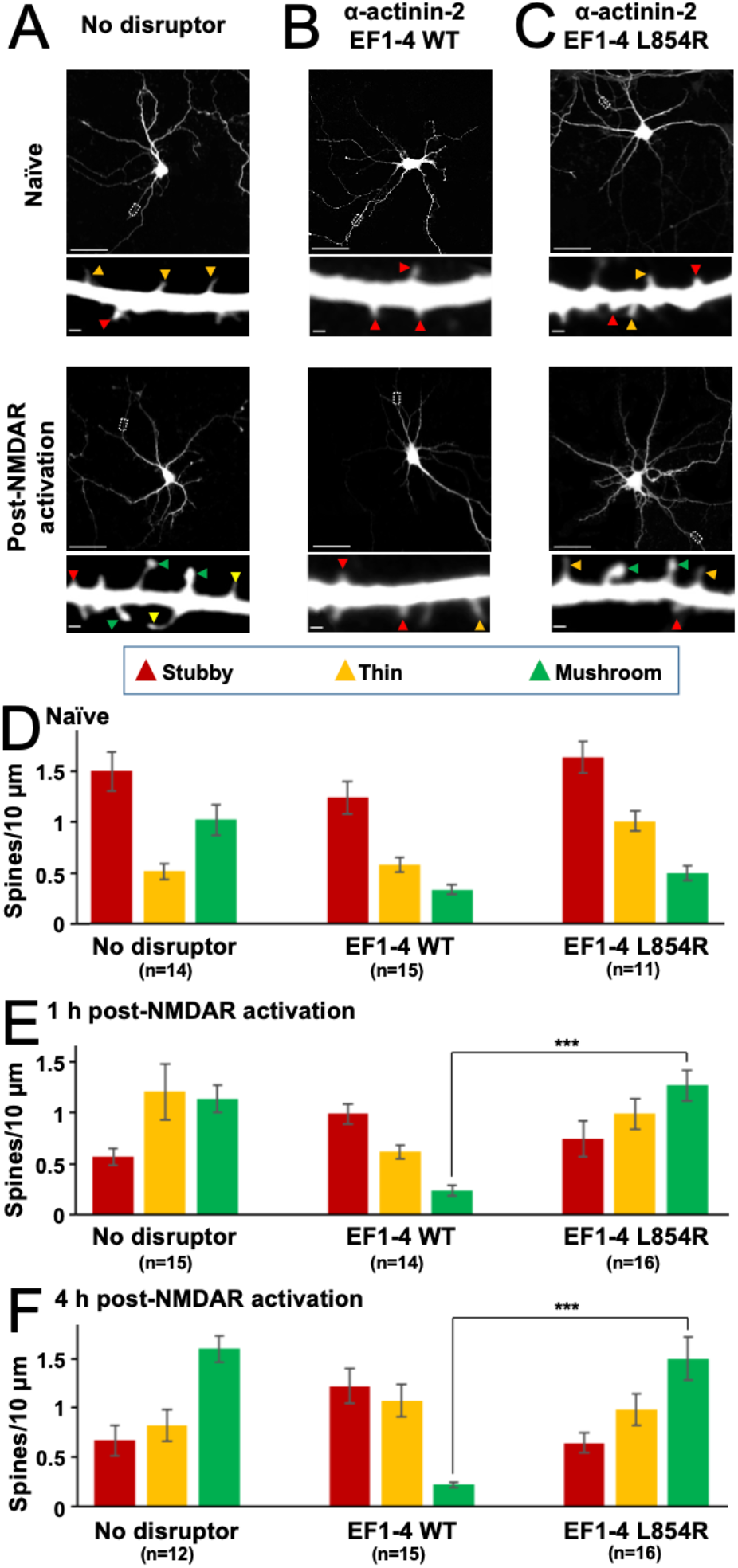
Effect of EF hand disruptors on spine dynamics following NMDAR activation. Panels A-C show GFP imaging of primary hippocampal neurons transfected with either GFP alone (A), or GFP in combination with α-actinin-2 EF1-4 WT (B) or L854R (C). For A-C, the upper rows show imaging of naïve synapses, and the lower rows shows imaging 4 hours after NMDAR activation. Stubby (red), thin (orange), and mushroom (green) type synapses are highlighted with arrows. Scale bars correspond to 20 *μm* (whole neuron images) and 1 *μm* (dendrite close-ups). Panels D-F show quantification of spine types across the three conditions either before NMDAR activation (D), 1 hour after NMDAR activation (E), or 4 hours after activation (F). Data are presented as mean ± SE spines per 10 *μm* dendritic length. The number of neurons analysed for each condition is shown in parentheses. Red, orange, and green bars indicate stubby, thin, and mushroom spine numbers respectively.

The L854R mutation falls within the fourth EF hand and disrupts α-actinin-2 binding to the CaMKII regulatory segment (Jalan-Sakrikar et al., 2012) thus serves as a negative control. PLA imaging confirmed that association with CaMKII was greatly reduced for the L854R variant (***Figure 2-figure supplement 1***). Prior to NMDAR activation, the total number of spines was approximately one third lower in neurons expressing the WT EF1-4 construct than in the other two conditions (***Figure 2-figure supplement 2***A). For all three conditions, the distribution of spine types was initially similar (***Figure 2***D), with stubby spines predominating at ~ 1.5 stubby spines per 10 μm dendrite prior to NMDAR activation. However, whereas NMDAR activation triggered a transition from stubby to mushroom type spines in neurons expressing GFP alone (***Figure 2***A) or GFP/EF1-4 L854R (***Figure 2***C), no such transformation occurred in neurons expressing the WT EF1-4 disruptor (***Figure 2***B).

Four hours after NMDAR activation (***Figure 2***F), stubby spines had decreased in abundance from 1.50±0.19 to 0.67±0.10/10 μm in GFP-only neurons (*p*=0.0013), and from 1.64±0.15 to 0.65±0.10/10 μm in neurons expressing L854R EF1-4 (*p*=7.2×10^-6^). Furthermore, mushroom spines accounted for approximately half of all spines four hours after NMDAR activation (***Figure 2***F). In contrast, mushroom spines did not accumulate in neurons expressing WT EF1-4 disruptor, such that after 4 hours there 7.2-fold fewer mushroom spines in these neurons than in GFP control neurons (*p*=3.43×10^-11^) and 6.8-fold fewer than in L854R EF1-4 neurons (*p*=4.81×10^-6^). The number of stubby spines also remained unchanged in WT EF1-4 neurons (1.24±0.16 before vs 1.22±0.18/10 μm four hours after NDMAR activation). The lack of plasticity in neurons expressing the WT EF1-4 disruptor was also reflected in analysis of changes in the average spine diameter upon NMDAR activation (***Figure 2-figure supplement 2***B). Mean spine diameter increased by 58.3 % (*p*=7.1×10^-8^) and 65.5 % (*p*=1.3×10^-4^) over four hours in GFP-only and L854R EF1-4 neurons, respectively whereas expression of the WT EF1-4 disruptor limited the increase to only 11 % (*p*=0.17). These results build on previous reports that siRNA-mediated knockdown of α-actinin-2 reduces mushroom-type spine formation following NMDAR activation (Hodges et al., 2014) by resolving a key role for the interface with CaMKII in this process. In sum, our PLA and spine imaging data indicate that α-actinin-2-CaMKII interactions accumulate upon induction of structural LTP via NMDAR activation, and that interaction between the two is necessary to support the formation of enlarged mushroom-type spines.

### The CaMKII kinase domain decreases affinity of α-actinin-2 for the regulatory segment

Previous studies have shown that EF hands 3 and 4 of α-actinin-2 (hereafter referred to as ‘EF3-4’) are sufficient for binding CaMKIIα via its regulatory segment (Jalan-Sakrikar et al., 2012; Robison, Bass, et al., 2005). Ca^2+^/CaM binds to CaMKII by employing its four EF hands to wrap around the regulatory segment. Since the segment is partially buried against the kinase domain in the inactive kinase (Chao et al., 2011; Rellos et al., 2010), Ca^2+^/CaM binds much more tightly to isolated regulatory segment than to constructs that include the kinase domain (Forest et al., 2008; Rellos et al., 2010). This is significant since factors that alter the accessibility of the regulatory segment, including T286 auto-phosphorylation (Singla et al., 2001) and association with NMDARs (Bayer et al., 2001), can trap CaM. For α-actinin-2, the assumption has been that EF3-4 is able to fully access the regulatory segment in inactive CaMKII since only two EF hands are involved in the interaction (Jalan-Sakrikar et al., 2012). However, our PLA data suggest that a mechanism exists for increasing CaMKIIα-actinin interactions following induction of LTP (***Figure 1***D). We therefore employed isothermal titration calorimetry (ITC) using purified proteins (***Figure 3-figure supplement 1***) to determine whether the kinase domain impedes access of α-actinin-2 to the regulatory segment in inactive CaMKII. We compared binding to a peptide corresponding to the regulatory segment (positions 294-315), and to a longer construct (1-315) that includes the kinase domain. An N-terminal thioredoxin (Trx) tag was fused at the N-terminus of the longer construct – where it would not be expected to affect interactions with the regulatory segment (Rellos et al., 2010) – to ensure that the protein remained soluble at high concentrations necessary for ITC (Costa et al., 2014). EF3-4 bound to the regulatory segment peptide (294-315) with a dissociation constant (K_d_) = 32±1 μM (***Figure 3***A). In contrast, it was not possible to determine a K_d_ for binding to the longer construct (1-315), with heat changes indistinguishable from background indicative of weaker binding (***Figure 3***B). Control experiments using CaM confirmed that, as expected, CaM binds more tightly to isolated regulatory segment (K_d_ = 11±1 nM, ***Figure 3***C) than to Trx-CaMKIIα 1-315 (K_d_ = 2.8±0.2 μM, ***Figure 3***D), consistent with previous studies (Rellos et al., 2010).

**Figure 3.**
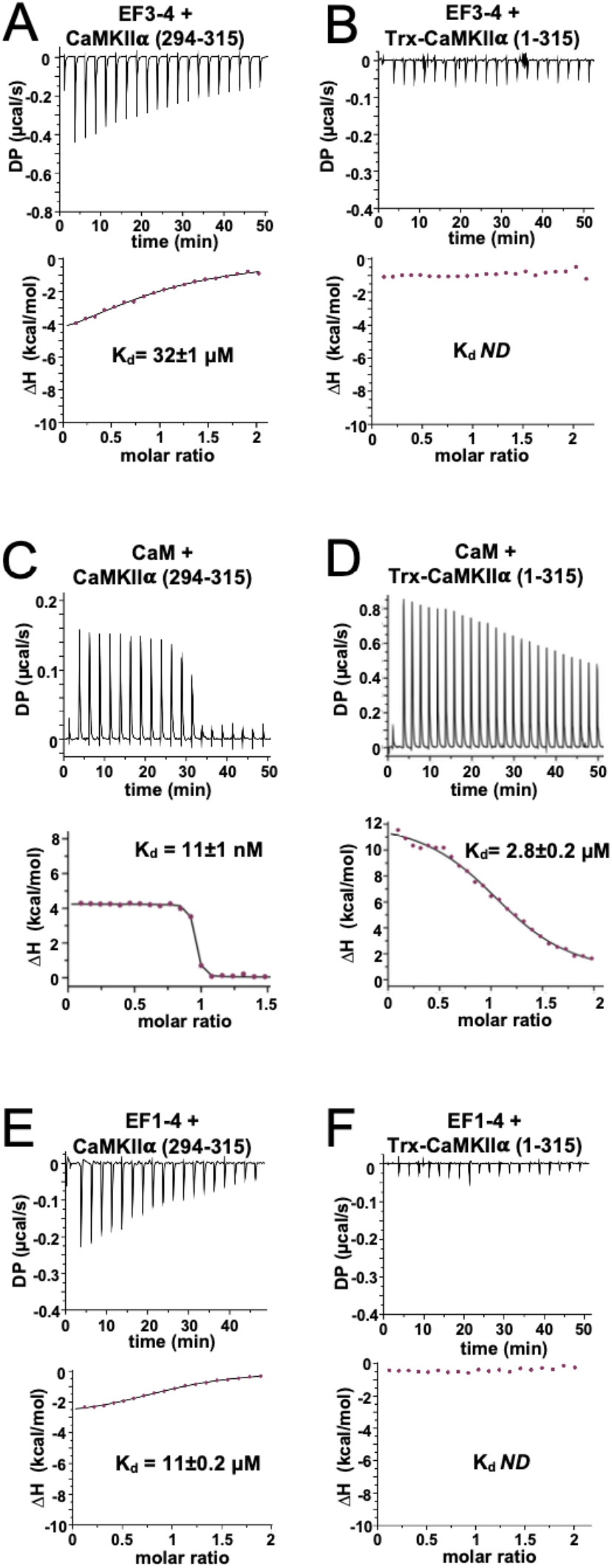
Isothermal titration calorimetry of interactions with the CaMKIIα regulatory segment. Representative isotherms showing binding of α-actinin-2 EF3-4 to (A) peptide corresponding to CaMKIIα regulatory segment (294-315) and (B) a construct (1-315) corresponding to the kinase and regulatory segment regions of CaMKIIα. Binding of CaM and α-actinin-2 EF1-4 to the same two CaMKII regions are shown in panel C&D and E&F, respectively. In all cases, the top sub-panels show the raw power output (μcal/s) per unit time; the bottom sub-panels show the integrated data including a line of best fit to a single site binding model. Stated K_d_ values are averages from experimental replicates. ND = not determined.

We next examined the notion that the first two EF hands of α-actinin-2 do not contribute to binding to the CaMKII regulatory segment (Jalan-Sakrikar et al., 2012; Robison, Bass, et al., 2005). Purified EF1-4 bound to the isolated regulatory segment (294-315) with K_d_=11±0.2 μM (***Figure 3***E) - a 3-fold lower concentration than EF3-4 alone (***Fig. 3*** A). This is similar to K_d_ = ~4 μM recorded for association of EF1-4 with peptide corresponding to titin Z repeat 7 (Grison et al., 2017). Like the shorter actinin construct (***Figure 3***B), no binding was detected between EF1-4 and Trx-CaMKIIα (1-315)(***Figure 3***F). Overall, our measurements show that – like CaM – α-actinin-2 is unable to fully access the CaMKII regulatory segment in the auto-inhibited enzyme. Furthermore, our data reveal that when the regulatory segment is fully accessible, all four EF hands of α-actinin-2 are required for the highest affinity binding. Full thermodynamic parameters obtained for all ITC measurements are shown in **Table 1**.

**Table 1.**
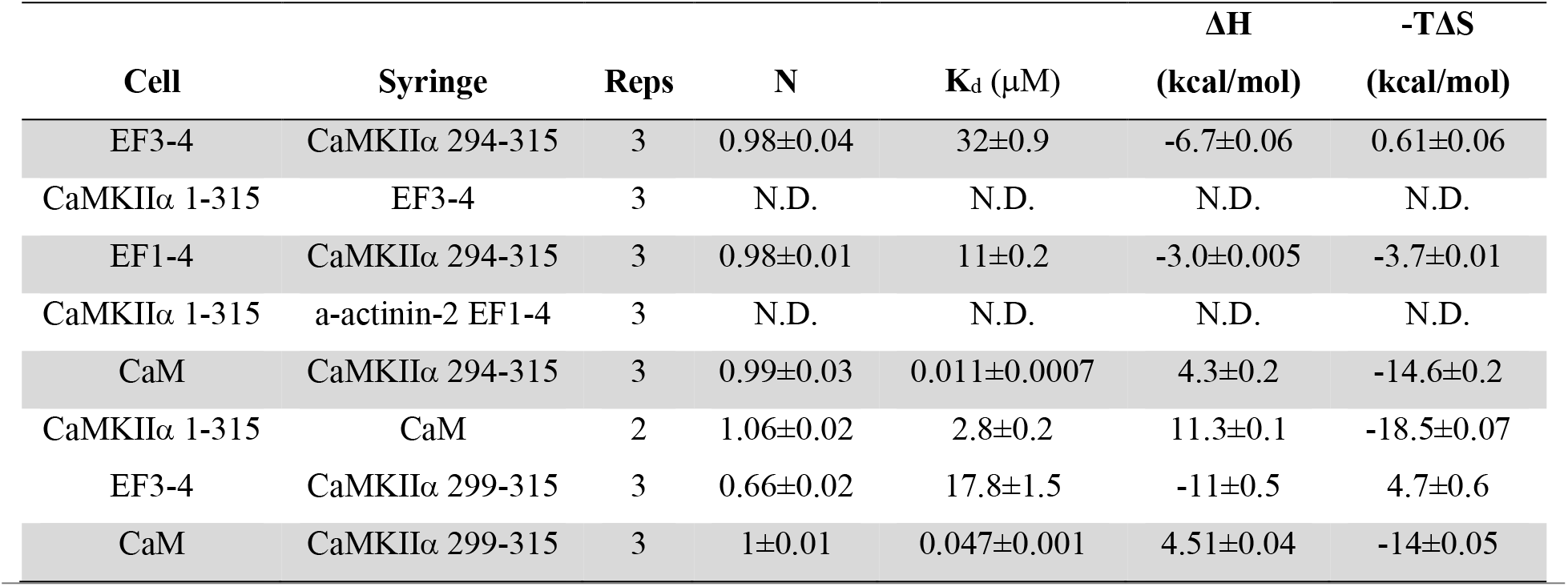
Thermodynamic parameters for interactions between α-actinin-2 and CaMKIIα constructs.

### Structure of the core α-actinin-2 – CaMKII interface

Previous structural models of α-actinin-2 – CaMKII interaction have assumed that the third and fourth EF hands of α-actinin-2 bind to the CaMKII regulatory segment in a similar mode to the third and fourth EF hands of Ca^2+^/calmodulin (Jalan-Sakrikar et al., 2012) in such a way that α-actinin-2 could fully access the regulatory segment in the inhibited kinase (Jalan-Sakrikar et al., 2012). However, our ITC measurements indicate that α-actinin-2 access to the regulatory segment is impeded in autoinhibited CaMKIIα (***Table 1***). Furthermore, PLA assays show increased association of the two proteins following chemical LTP protocols that activate CaMKII (***Figure 1***). To establish a structural basis for these findings, we crystallized α-actinin-2 EF3-4 in complex with a peptide corresponding to the CaMKIIα regulatory segment (positions 294-315).

We solved the crystal structure at 1.28 Å resolution (***Figure 4-figure supplement 1***A) using x-ray diffraction with phasing through single-wavelength anomalous diffraction of native sulfur atoms (***Supplementary Table 1***). The asymmetric unit contains two copies of the complex (***Figure 4-figure supplement 1***B) – the two copies are highly similar (RMSD 0.202 Å for all atoms, ***Figure 4-figure supplement 1***C). Our analysis focuses on the first copy of the complex (chains A & B in PDB ID 6TS3) for which it was possible to position the full EF3-4 region (824-894) and positions 294-313 of the CaMKIIα regulatory segment in electron density.

In the complex, the CaMKII regulatory segment (blue, ***Figure 4***A) forms an amphipathic helix with four hydrophobic residues (L299, I303, M307, F313) aligned and engaged in van der Waals interactions with α-actinin-2 EF3-4 (orange, ***Figure 4***A). M307_CaMKII_ is buried closest to the centre of the EF3-4 domain, where it contacts the sidechains of EF3-4 residues F835 and L854. This binding mode is consistent with the reduction in PLA puncta that we observed with the EF3-4 mutation L854R (***Figure 2-figure supplement 1***) and with earlier pull-down experiments (Jalan-Sakrikar et al., 2012). α-actinin-2 mutations S834R or Y861R have also been found to prevent association with CaMKII (Jalan-Sakrikar et al., 2012), and both of these residues also directly contact the regulatory segment in the crystal structure. The most N-terminal amino acid in the regulatory segment to engage directly with EF3-4 is L299_CaMKII_, which interacts with Y861_EF3-4_ (***Figure 4**A*): unexpectedly, positions 294-298 are solvent exposed. At the other end of the regulatory segment, the benzene ring of F313_CaMKII_ packs against the sidechain of I837_EF3-4_ (***Figure 4***B). Deletion studies have found that the last 9 amino acids (886-894) of α-actinin-2 are necessary for binding CaMKII (Robison, Bass, et al., 2005). This region includes Y889_EF3-4,_ which engages in van der Waals interactions with L304_CaMKII_ through its benzene ring and H-bonds with the backbone oxygen of K300_CaMKII_. The last four amino acids of α-actinin-2 (‘ESDL’) form a potential ligand for PDZ domains in proteins including densin-180 (Robison, Bass, et al., 2005; Walikonis et al., 2001). In the structure, G890_EF3-4_ reorients the polypeptide chain away from the interface with CaMKII such that this PDZ ligand is accessible (***Figure 4***B). The CaMKII regulatory segment is subject to negative feedback auto-phosphorylation at T305 and T306, with phosphorylation at either site preventing further activation by CaM (Hanson & Schulman, 1992; Patton et al., 1990). Mutational analyses have previously shown that phosphorylation at T306 but not T305 reduces binding to α-actinin-2 (Jalan-Sakrikar et al., 2012). This tallies with the crystal structure, in which T305 projects away from the interface with EF3-4 whereas the hydroxyl group of T306 engages in a H-bond network that includes Gln858_EF3-4_, and its Cg atom is packed against P885_EF3-4_ (***Figure 4-figure supplement 1***D). The EF3-4 region adopts a similar conformation in complex with the CaMKII regulatory segment as in the crystal structure of full-length apo α-actinin-2 (Almeida Ribeiro Jr. et al., 2014) with 1.168 Å RMSD for all atoms. The CaMKIIα regulatory segment (blue, ***Figure 4***C) occupies the same position as the neck region of α-actinin-2 (grey, ***Figure 4***C). In neurons, association of the actin-binding domain of α-actinin-2 with actin filaments, and binding of PIP2 phospholipids in the vicinity of the neck region, are thought to ensure that EF3-4 is available for interaction with CaMKII (Almeida Ribeiro Jr. et al., 2014; Hell, 2014). The neck region residue Ile269 occupies the equivalent position to Met307_CaMKII_ (***Figure 4***A), although the polypeptides run in opposite directions across the EF3-4 region.

**Figure 4.**
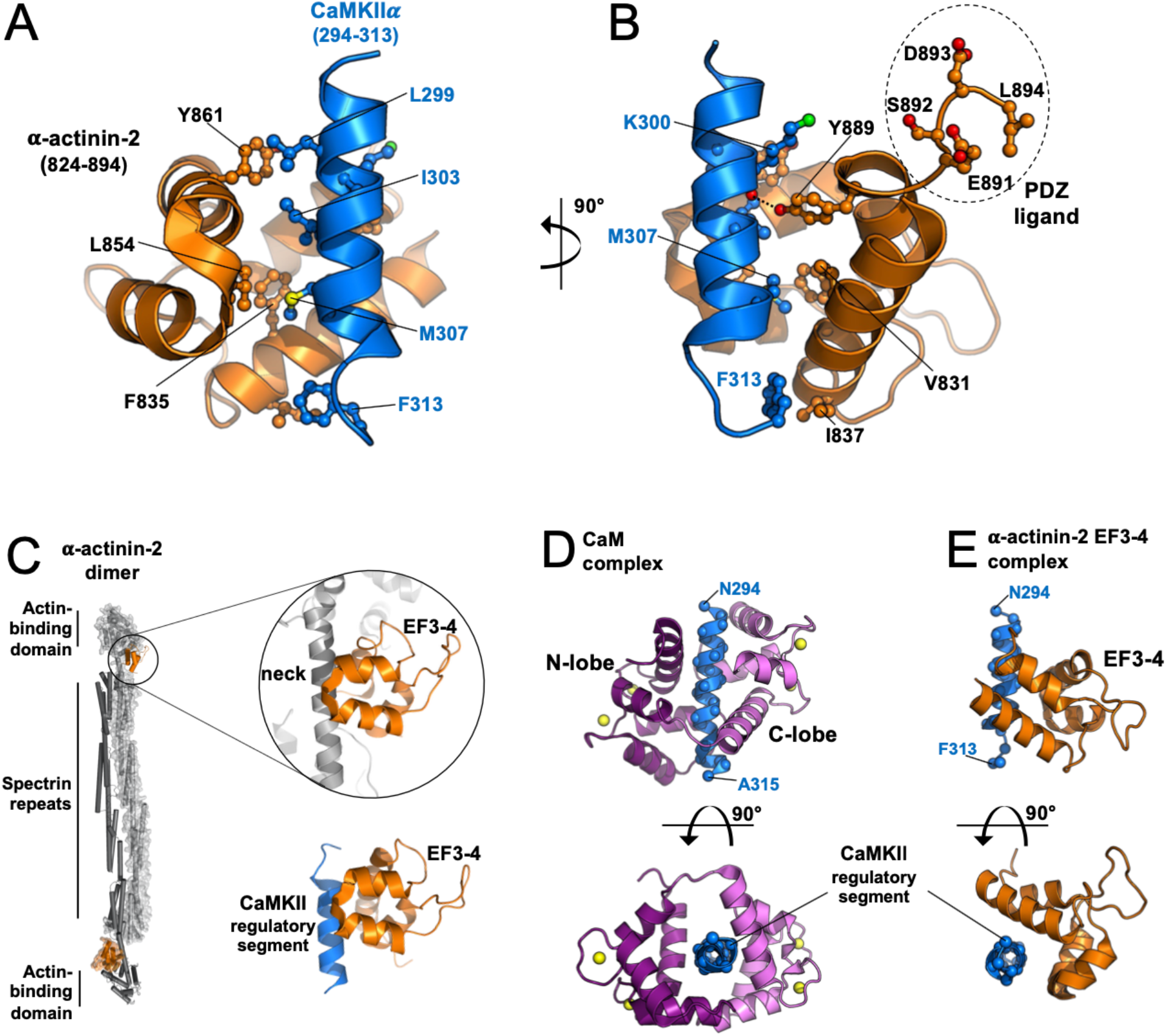
Structure of the core α-actinin-2 - CaMKIIα interface. Panels (A) and (B) show two views of the complex between α-actinin-2 EF3-4 (orange) and a peptide corresponding to the CaMKIIα regulatory segment (blue). The C-terminal tetrapeptide that is a ligand for PDZ domain-containing proteins including densin-180 is highlighted. (C) Comparison of α-actinin-2 EF3-4 domain association with the neck region (grey) in α-actinin-2 dimers (1H8B) and the CaMKIIα regulatory segment (6TS3). The structures were aligned through the EF3-4 domain. Panels (D) and (E) show two views comparing CaM (D) and α-actinin-2 EF3-4 (E) association with the CaMKIIα regulatory segment. The structures were aligned through the regulatory segment. For the CaM complex (2WEL), the N-lobe is coloured dark purple; the C-lobe is violet.

Previous modelling of the CaMKII-actinin interaction has been built on the assumption that α-actinin-2 EF3-4 binds to CaMKII using the same binding mode as the CaM C-lobe (Jalan-Sakrikar et al., 2012). However, comparison to the crystal structure of CaMKIIδ (11-315) in complex with Ca^2+^/CaM (Rellos et al., 2010) shows that the binding modes are distinct. Ca^2+^/CaM fully encompasses the regulatory segment (whose sequence is identical between the a and δ isoforms) with the C-lobe (violet, ***Figure 4***D) mediating most interactions to the hydrophobic face of the helix. The N-lobe (deep purple) is responsible for the bulk of H-bonding to the helical side that is solvent exposed in complex with α-actinin-2 EF3-4 (***Figure 4***E). The centre of mass of EF3-4 is rotated by ~ 50 ° relative to the CaM C-lobe when viewed along the central axis of the regulatory segment (lower panels, ***Figure 4***D & E). In addition, the CaM C-lobe engages the regulatory segment approximately one helical turn further to its N-terminus than α-actinin-2 EF3-4, including interactions with positions A295 and K298. The conformation of the regulatory segment itself also differs between the two complexes. In complex with α-actinin-2 EF3-4, the helical structure breaks down at the C-terminal end to re-orient F313 for interaction with I837_EF3-4_ (light blue, ***Figure 4-figure supplement 1***E). In complex with Ca^2+^/CaM, alpha-helicity is maintained for the full length of the regulatory segment, which directs F313 in the opposite direction for interaction with the CaM N-lobe (dark blue, ***Figure 4-figure supplement 1***E). Since CaMKIIα positions 294-298 are solvent exposed in the complex with α-actinin-2 EF3-4 but not CaM, we performed further ITC measurements with a truncated regulatory segment peptide (299-315) to corroborate the binding mode observed in the crystal structure. α-actinin-2 EF3-4 bound CaMKIIα 299-315 peptide with K_d_ = 17.8±1.5 μM (***Figure 4-figure supplement 2***A) – comparable to its affinity for CaMKIIα 294-315 (K_d_ = 32±1). Consistent with the crystal structures, CaM associated with CaMKIIα (299-315) with K_d_ = 47±1 nM (***Figure 4-figure supplement 2***B) – approximately 5-fold higher than for the longer peptide (K_d_ =11±1 nM. Overall, our structural data show how α-actinin-2 employs a unique binding mode to interact with the CaMKII regulatory segment.

### Occlusion of the regulatory segment to α-actinin-2 can be released by association with GluN2B

The reduced affinity of EF3-4/EF1-4 for the CaMKII regulatory segment in constructs that include the kinase domain (**Table 1**) suggests that the kinase domain impedes α-actinin-2 access to the regulatory segment. To understand the structural basis of this effect, we superimposed the EF3-4 – regulatory segment complex structure on previously-determined crystal structures of auto-inhibited CaMKII. ***Figure 5***A & B show superimposition onto the structure of CaMKIIα (13-302) bound to the inhibitor indirubin (Rellos et al., 2010). In this structure L299 is the last visible residue in CaMKII (Rellos et al., 2010). The common residues 294-299 align closely with RMSD = 0.2 Å for backbone carbon and oxygen atoms (***Figure 5-figure supplement 1**A*). The superimposition shows that – without reorientation or partial release of the regulatory segment from the kinase domain – both termini of the EF3-4 region will clash with the kinase domain (***Figure 5**A* & B). Steric incompatibility of the kinase domain and EF3-4 region is most evident in the vicinity of the kinase domain aG helix (***Figure 5***C), which is located in the equivalent position to the last 5 amino acids of α-actinin-2. We also superimposed the EF3-4 – regulatory segment complex onto the crystal structure of a chimeric form of full-length CaMKII, which incorporates WT sequence for CaMKIIα up to position T305 (Chao et al., 2011) (***Figure 5-figure supplement 1***B & C). The two structures were aligned on the basis of the common positions 294-305. In this case, steric clashing was more pronounced since the regulatory segment kinks towards the kinase domain in this case (***Figure 5-figure supplement 1***C).

**Figure 5.**
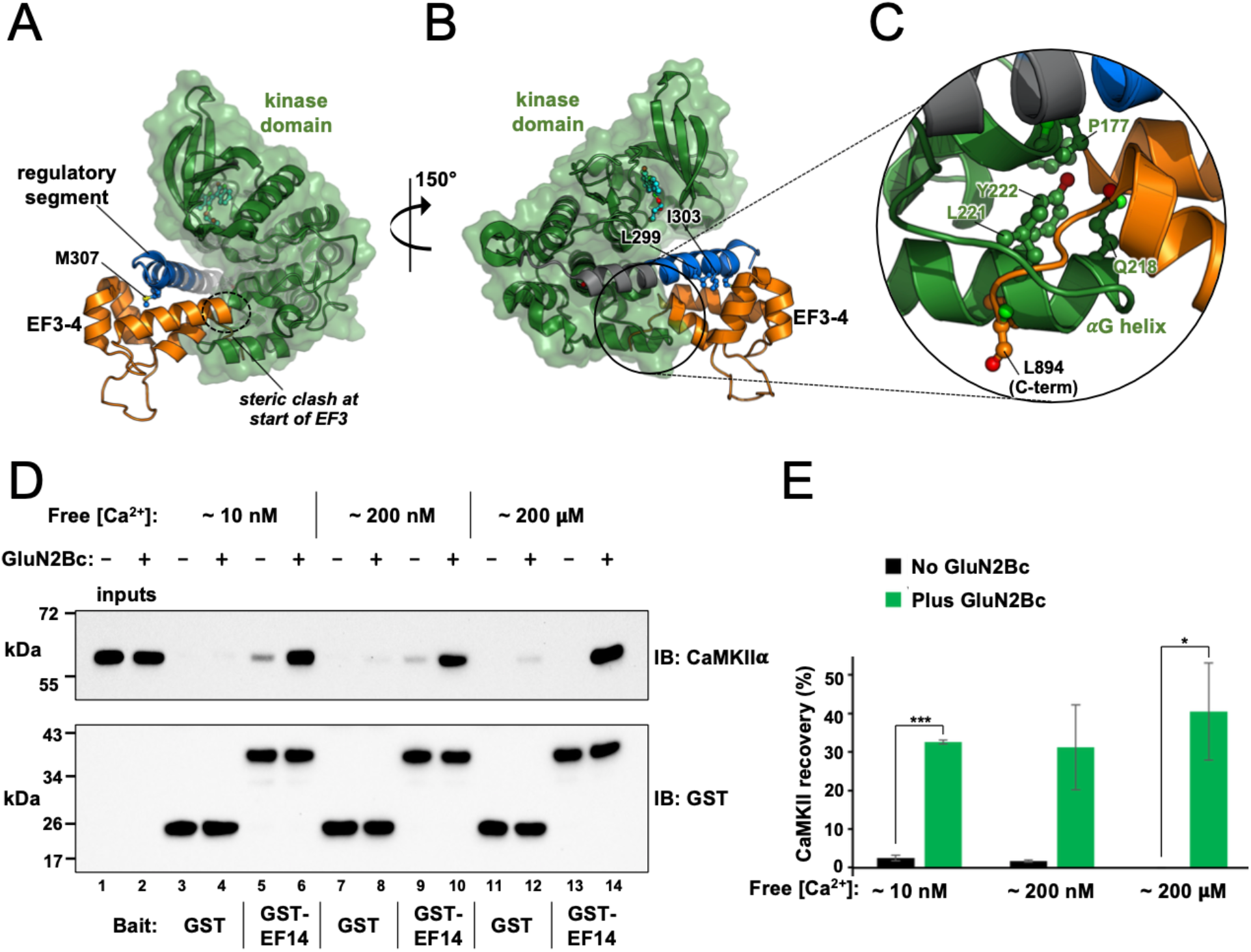
Effect of GluN2B on access to the regulatory segment. Panels (A) and (B) show two views of the superimposition of the α-actinin-2 – regulatory segment complex (6TS3) onto the structure of CaMKIIα 1-299 (2VZ6). The structures were aligned using positions 294-299 of the regulatory segment. Regions of steric incompatibility between the kinase (green) and EF3-4 domain (orange) are highlighted. Panel C shows a close-up highlighting clashing in the vicinity of the CaMKII αG helix. (D) Pull-down of purified CaMKIIα with magnetic beads charged with either GST or GST-EF1-4. CaMKII pull-down was compared +/− GluN2Bc fragment, and at different final free Ca^2+^ concentrations, as indicated. CaMKIIα and GST/GST-EF14 were detected by anti-CaMKIIα (upper) and anti-GST (lower) immunoblots (IBs). (E) Densitometry for pull-down experiments shown in the preceding panel showing CaMKII recovery at each free Ca^2+^ concentration either with (green) or without (black) GluN2Bc (n=3 for all conditions).

We found that α-actinin-2 association with CaMKII was markedly increased in dendritic spines four hours after chemical induction of LTP by NMDAR activation (**Figure 1**). Since our structural and calorimetry data show that the kinase domain occludes α-actinin-2 access to the regulatory segment in autoinhibited CaMKII, the mechanism underlying this increase in association is likely to involve regulatory segment release. Existing knowledge of CaMKII regulation during LTP suggests two possibilities: auto-phosphorylation at T286, and formation of NMDAR-CaMKII complexes (Yasuda et al., 2022). During the initial phase of LTP induction, CaMKII auto-phosphorylates at T286 to generate an autonomously active form with partial activity of ~ 20 % (Barcomb et al., 2014). The pT286 modification is likely to partially disengage the regulatory segment from the kinase domain, however this modification does not endure for more than ~ 10 s following the induction of LTP (Chang et al., 2017; Yasuda et al., 2022). A more logical possibility to explain increased actinin-CaMKII association hours after LTP induction is the formation of NMDAR-CaMKII complexes that endure hours following CaMKII activation (Bayer et al., 2001; Yasuda et al., 2022). CaMKII binds tightly to a site centred on S1303 in the GluN2B C-terminal tail (Omkumar et al., 1996; Strack, McNeill, et al., 2000), and recent crystallographic work shows how this sequence wraps around the kinase domain using a binding mode that necessitates full displacement of the regulatory segment from the kinase domain (Ozden et al., 2022). To investigate the possibility that GluN2B supports CaMKIIα binding to α-actinin-2, we compared pull-down of full-length CaMKIIα with magnetic glutathione beads charged with either GST or GST-EF1-4 and determined the effect of including a purified fragment of the GluN2B tail spanning positions 1260-1492 (***Figure 3-figure supplement 1***E) – hereafter referred to as ‘GluN2Bc’. In all cases, an initial charging step was included in 2 mM CaCl_2_ to enable CaMKII-GluN2B association prior to addition of EGTA and incubation with the charged magnetic beads. Reaction mixtures were supplemented with 2 % BSA, and incubated with the magnetic beads for only 20 minutes to reduce basal pull-down in the absence of GluN2Bc. We also compared CaMKII pull-down with three final EGTA concentrations: 10, 2.5, or 1.8 mM corresponding to final approximate free Ca^2+^ concentrations of 15 nM, 0.4 μM, and 200 μM (***Figure 5***D). CaMKII recovery without GluN2B was reduced from 2.5±0.8 % to 0±0.1 % (*p*=0.027, ***Figure 5***E) in the absence of GluN2B moving from low to high final Ca^2+^ (lanes 5 & 13, ***Figure 5***D), consistent with previous reports that Ca^2+^/CaM alone outcompetes α-actinin-2 for binding to CaMKII (Robison, Bartlett, et al., 2005). Some association without GluN2Bc at low Ca^2+^ levels is consistent with our PLA imaging (***Figure 1***D) and with the original identification of the actinin-CaMKII interaction by the yeast two-hybrid method (Walikonis et al., 2001) and it should be noted that this baseline interaction is more pronounced under less stringent binding conditions (Jalan-Sakrikar et al., 2012).

Addition of GluN2Bc led to striking increases in CaMKII recovery irrespective of EGTA concentration. At ~15 nM free Ca^2+^, GluN2Bc increased recovery from from 2.5±0.8 % to 33±0.5 % (lanes 5&6, ***Figure 5***B, *p*=5.0×10^-6^). At 0.4 μM Ca^2+^, recovery increased from 1.7±0.3% to 31±11 % (lanes 9&10, *p*=0.055). Surprisingly, the effect was maintained at 200 μM free Ca^2+^, with GluN2Bc increasing recovery from 0±0.1 % to 40±13 % (lanes 13&14, ***Figure 5***D, *p*=0.032). The effects of GluN2Bc on CaMKII recovery are summarised in ***Figure 5***E. Overall, these experiments indicate that association of CaMKII with GluN2B subunits following LTP is a plausible mechanism to enable increased interaction between the kinase and α-actinin-2 in structural LTP.

## Discussion

We propose a revised mechanism for spine enlargement in structural LTP (**Figure 6**) based on our findings. In naïve spines, the regulatory segment of CaMKII is sequestered by its kinase domain (***Figure 6***A) within inactive dodecamers, ensuring only limited association with α-actinin-2 in the ground state (***Figure 1***D). Upon induction of LTP by Ca^2+^ influx through NMDARs, activated Ca^2+^/CaM binds to the regulatory segment of CaMKII which exposes the substrate binding groove of the kinase domain (***Figure 6***B). The activated kinase phosphorylates key substrates including AMPA receptors and forms a highly-stable complex with NMDARs by interaction with a substrate binding site within the C-tail of GluN2B subunits. The duration of CaMKII activation is prolonged beyond the duration of Ca^2+^ elevation by auto-phosphorylation at T286, although pT286 autonomy is reversed by phosphatases including protein phosphatase 1 within ~ 10 s (Yasuda et al., 2022). In complex with NMDARs, the substrate-binding groove of the CaMKII kinase domain is occupied (Ozden et al., 2022), which leaves the regulatory segment accessible to interact with α-actinin-2 (***Figure 6***C). In this way, α-actinin-2 is enriched in dendritic spine heads where it supports spine head enlargement through its ability to crosslink actin filaments via its N-terminal actin-binding domain (***Figure 6***C). This updated mechanism fits with an influential ‘synaptic tagging’ theory (Sanhueza & Lisman, 2013), whereby long-lasting CaMKII-NMDAR complexes formed during LTP induction serve as primers for more global structural changes in dendritic spines that occur after the initial Ca^2+^ signal has subsided.

**Figure 6.**
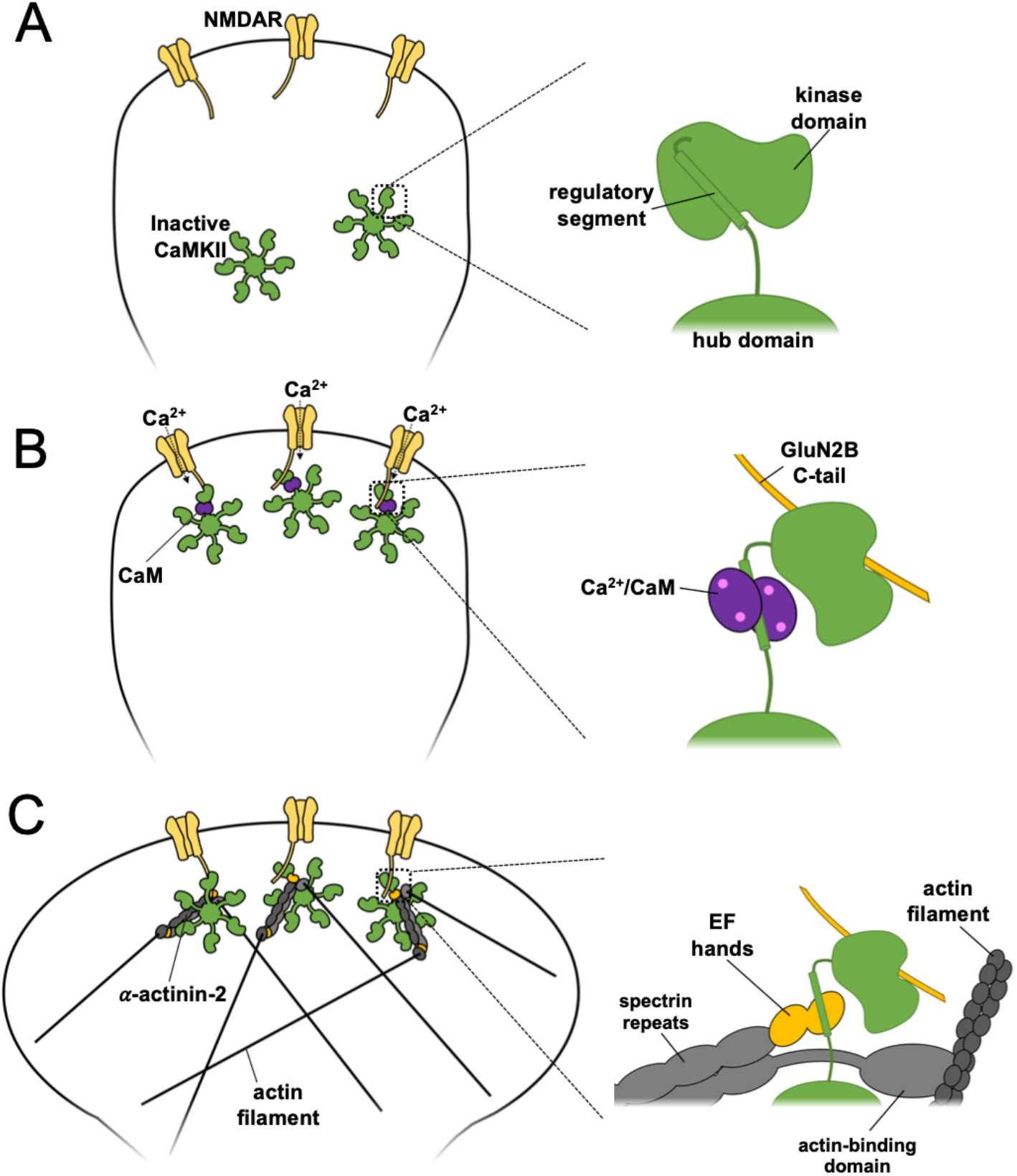
Model of actinin-CaMKII dynamics underlying structural LTP. A three-stage model is presented with close-up illustrations of protein interactions involving CaMKII on the right for each stage. (A) In naïve synapses prior to Ca^2+^ entry, the CaMKII regulatory segment associates with the kinase domain and is largely inaccessible to α-actinin-2. (B) Ca^2+^ influx through NMDARs triggers binding of Ca^2+^/CaM (purple) to the regulatory segment, enabling docking of the kinase domain to the C-terminal tail of GluN2B (gold) subunits. (C) Following return of spine [Ca^2+^] to baseline levels, Ca^2+^/CaM dissociates whereas CaMKII-NMDAR interactions persist. This combination enables *α*-actinin-2 to dock to the kinase domain, thereby linking the kinase to the actin cytoskeleton in support of spine enlargement.

The simplified model presented in ***Figure 6*** is broadly consistent with many additional protein-protein interactions involving CaMKII and α-actinins that are known to contribute to the positioning of the two proteins in dendritic spines. Densin-180 includes a C-terminal PDZ domain that binds to the PDZ ligand at the C-terminus of α-actinin-2 (Walikonis et al., 2001), and a motif ~100-150 amino acids from its C-terminus that associates with the CaMKII hub domain (Strack, Robison, et al., 2000). In the crystal structure (***Figure 4***B), the PDZ ligand at the C-terminus of EF3-4 is available for interaction which suggests that densin-180 could form a tripartite complex with CaMKII and α-actinin including a single CaMKII protomer. α-actinin-2 is also known to interact with other NMDAR subunits, and PSD-95. α-actinin-2 binds to the C-tails of GluN1 subunits (Wyszynski et al., 1997) through its central spectrin repeats (***Figure 1***A), which could be compatible with binding to CaMKII associated with GluN2B subunits. α-actinins also associate with the N-terminal 13 amino acids of PSD-95, which underlie their role in spine formation (Matt et al., 2018), further supporting a role for the actin-crosslinking protein in the PSD. Accurately conceptualising interactions between proteins in the PSD is complicated by the potential for cooperative interactions since many of the relevant proteins are oligomeric. For example, it is not clear whether EF3-4 domains at either end of an α-actinin-2 dimer could simultaneously bind to two regulatory segments within the same CaMKII dodecamer (Penny & Gold, 2018). Furthermore, NMDAR receptors themselves are spaced at regular intervals of ~ 30 nm in the PSD (Chen et al., 2008) – a similar scale to the width of CaMKII dodecamers (~ 25 nm) and the length of α-actinin-2 dimers (~ 28 nm)(Almeida Ribeiro Jr. et al., 2014; Myers et al., 2017) – raising the possibility of cooperative interactions spanning multiple receptors. Developments in techniques for *in situ* imaging including cryo-electron tomography (van den Hoek et al., 2022) will be required to resolve the extent to which cooperative interactions support the structure of potentiated dendritic spines.

Our imaging experiments fit with previous reports that α-actinin-2 knockdown prevents formation of mushroom-type dendritic spines (Hodges et al., 2014). α-actinin-2 is best known for its role at the Z disk in cardiac, striated, and smooth muscle cells where it organises the lattice structure of the contractile apparatus through crosslinking actin filaments and binding Titin through the EF3-4 region (Sjoblom et al., 2008). Actin is not visible in equivalent arrays with regular spacing in dendritic spines (Burette et al., 2012). In this location, α-actinins likely operate through a mechanism more analogous to their role during cytokinesis where they accumulate actin filaments at the cleavage furrow without generating a regular lattice structure (Mukhina et al., 2007). An aspect of CaMKII regulation of spine architecture that we did not consider in this study is the role of CaMKIIβ subunits, that bind directly to the actin cytoskeleton through elements within their regulatory segment (O’Leary et al., 2006; Shen et al., 1998). Only a fraction of CaMKII dodecamers contain β subunits (Miller & Kennedy, 1985) but within these assemblies it is possible that tripartite complexes assemble including CaMKII, α-actinin and F-actin. In our summary model (***Figure 6***), we suggest that docking to NMDARs is key for enabling α-actinin-2 to access the CaMKII regulatory segment but potentially other postsynaptic proteins that stably associate with CaMKII by occupying the substrate-binding groove of the kinase domain could also support activity-dependent increases in CaMKII-actinin association. There is potential for therapeutic inhibition of CaMKII (Pellicena & Schulman, 2014), and several leading peptide inhibitors of the kinase constitute pseudosubstrate sequences that occupy the substrate binding groove (Reyes Gaido et al., 2022). Our work suggests that developers of such inhibitors should be mindful of the potential for unexpected gain-of-function effects in inhibitors that increase access to the regulatory segment. Overall, our study illuminates the remarkable sophistication of regulatory processes that enable a single kinase – CaMKII – to play such a central role in the regulation of synaptic strength.

## Methods & Materials

### Key Resources Table

| Reagent type<br>(species) or<br>resource | Designation | Source or<br>reference | Identifiers | Additional<br>information |
| --- | --- | --- | --- | --- |
| strain, strain<br>background | TOP10<br>chemically<br>competent *E.*<br>*coli* | Life<br>Technologi<br>es | Cat# C404003 |  |
| strain, strain<br>background | *E. coli* BL21<br>(DE3) | Thermo<br>Fisher<br>Scientific | Cat# EC0114 |  |
| strain, strain<br>background | *E. coli* BL21<br>Tuner (DE3)<br>pLysS | Merck | Cat# 70624 |  |
| strain, strain<br>background | *E. coli* BL21<br>Star (DE3) | Thermo<br>Fisher<br>Scientific | Cat# C601003 |  |
| cell line | HEK293T<br>(human) | ATCC | Cat# CRL-3216 | Mycoplasma-tested |
| biological<br>sample | Sprague<br>Dawley rat | UCL<br>breeding<br>colony | Not applicable |  |
| antibody | Rabbit anti-GFP | Sigma<br>Aldrich | Cat# SAB4301138;<br>RRID:AB_2750576 | 10 µg/mL |
| antibody | Goat anti-FLAG | Novus | Cat# NB600-344;<br>RRID:AB_10000565 | 5 µg/mL |
| antibody | Rabbit anti-GST | Merck | Cat# G7781;<br>RRID:AB_259965 | 0.2 µg/mL |
| antibody | Mouse anti-CaMKIIα | Santa Cruz Biotechnology | Cat# sc-32288;<br>RRID:AB_626787 | 0.2 µg/mL for IB; 0.5 µg/mL for ICC |
| antibody | Goat anti-rabbit HRP-linked secondary antibody | Cell Signalling Technology | Cat # 7074S;<br>RRID:AB_2099233 | 0.5 µg/mL |
| antibody | Goat anti-mouse poly-HRP secondary antibody | Thermo Fisher Scientific | Cat# 32230;<br>RRID:AB_1965958 | 0.5 µg/mL |
| antibody | goat anti-rabbit Alexa Fluor-405 | Thermo Fisher Scientific | Cat# A-31556;<br>RRID:AB_221605 | 4 µg/mL |
| antibody | Duolink In Situ PLA Probe Anti-Mouse MINUS | Merck | Cat# DUO92004 |  |
| antibody | Duolink In Situ PLA Probe Anti-Goat PLUS | Merck | Cat# DUO92003 |  |
| recombinant DNA reagent | pIRES2-EGFP | Clontech | Cat# 6029-1 |  |
| recombinant DNA reagent | pIRES2-EGFP-FLAG-α-actinin-2 construct series (1-894;<br>1-748; 747-894; 747-890;<br>1-890; 747-894 L854R) | This study | Not applicable |  |
| recombinant<br>DNA reagent | pcDNA3.1-6His-CaMKIIα | This study | Not applicable | |
| recombinant<br>DNA reagent | pET28-6His-GST-Tev-<br>actinin-2 EF3-4 (827-894) | This study | Not applicable |  |
| recombinant<br>DNA reagent | pNH-TrxT-6His-Trx-Tev-CaMKIIα (1-315) | This study | Not applicable | |
| recombinant<br>DNA reagent | pGEX6P1-α-<br>actinin-2 EF1-4 (747-894) | This study | Not applicable | |
| recombinant<br>DNA reagent | pGEX6P1-GluN2B<br>(1260-1492) | This study | Not applicable |  |
| recombinant<br>DNA reagent | pET28-CaM<br>(untagged) | (Patel et al.,<br>2017) | Not applicable |  |
| recombinant<br>DNA reagent | pCI-EGFP-NR2b | Andres<br>Barria &<br>Robert<br>Malinow/A<br>ddgene | RRID:Addgene_45447 |  |
| recombinant DNA reagent | pNH-TrxT | Opher Gileadi/Ad dgene | RRID:Addgene_26106 |  |
| recombinant DNA reagent | pET3d-6His-α-actinin-2 | Kristina Djinović-Carugo (Almeida Ribeiro Jr. et al., 2014). | Not applicable | |
| chemical compound, drug | Lipofectamine 2000 | Thermo Fisher Scientific | Cat# 11668019 |  |
| chemical compound, drug | Polyethylenimine (linear, MW25000, transfection grade | Polysciences | Cat# 23966 |  |
| chemical compound, drug | Duolink In Situ Detection Reagents Red | Merck | Cat# DUO92008 |  |
| chemical compound, drug | Pierce Glutathione Magnetic Agarose Beads | Thermo Fisher Scientific | Cat# 78601 |  |
| chemical compound, drug | DMEM, high glucose, pyruvate | Thermo Fisher Scientific | Cat # 41966029 |  |
| chemical compound, drug | Trypsin | Thermo Fisher Scientific | Cat# 25300054 |  |
| chemical compound, drug | Penicillin/Streptomycin | Thermo Fisher Scientific | Cat# 15140122 |  |
| chemical compound, drug | GlutaMAX | Thermo Fisher Scientific | Cat# 35050061 |  |
| chemical compound, drug | DPBS, no calcium, no magnesium | Thermo Fisher Scientific | Cat# 14190144 |  |
| chemical compound, drug | Neurobasal-A medium | Thermo Fisher Scientific | Cat# 10888022 |  |
| chemical compound, drug | B27 supplement | Gibco | Cat# 17504044 |  |
| chemical compound, drug | Poly-L-Lysine | Sigma Aldrich | Cat# P2636 |  |
| chemical compound, drug | cOmplete, Mini, EDTA-free Protease Inhibitor Cocktail | Roche | Cat# 11836170001 |  |
| software, algorithm | ImageJ (version 1.52) | NIH | RRID:SCR_003070 |  |
| software, algorithm | NeuronStudio | (Rodriguez et al., 2008) | https://icahn.mssm.edu/; RRID:SCR_013798 |  |

#### Protein expression and purification

The EF3-4 (827-894) region of human α-actinin-2 was cloned into a modified pET28 vector using primers EF34_F & _R for expression with a Tev-cleavable N-terminal 6His-GST tag. The construct was transformed into Rosetta plysS *E. coli*, and expression was induced with 1 mM IPTG after the cells had reached an OD_600nm_ of ~ 0.7 in LB. Cells were harvested after overnight incubation at 20 °C, resuspended in Ni-NTA buffer A (25 mM Tris pH 8, 500 mM NaCl, 1 mM Benzamidine, 30 mM imidazole) supplemented with 1 cOmplete protease inhibitor tablet/100ml and 0.1 mg/mL lysozyme, then clarified by centrifugation at 40,000 x *g*. The clarified lysate was incubated with Ni-NTA agarose beads (Qiagen) for 1.5 hours. Following incubation, beads were washed and eluted (500 mM NaCl, 25 mM Tris pH 7.5, 1 mM Benzamidine, 300 mM imidazole). The resulting eluate was buffer exchanged into Glutathione Sepharose binding buffer (25 mM HEPES pH 7.5, 500 mM NaCl, 1 mM DTT, 0.5 mM EDTA) and incubated with glutathione Sepharose 4B beads for 3 hours, washed and cleaved overnight at 4 °C with Tev protease. Following overnight cleavage, samples were further purified by size-exclusion chromatography with a HiLoad 16/600 Superdex 75 column equilibrated in 20 mM Na HEPES pH 7.5, 0.15 M NaCl, and 1 mM DTT.

Coding sequences for rat GluN2B (1266-1482) and α-actinin-2 EF1-4 (747-894) were cloned into pGEX-6P1 using primers BamHI_GluN2B_1260/GluN2B_Term_EcoRI and EcoRI_actinin_747/actinin_Term_NotI, respectively. GluN2Bc was expressed in Tuner (DE3) *E.coli* whereas EF1-4 was expressed in Rosetta plysS *E. coli*. In both cases, bacteria were harvested following overnight growth in auto-inducing media (AIM)(Studier, 2005) at 37 °C. The two protein fragments were purified in the same way. Bacteria were first lysed by sonication in lysis buffer (25 mM HEPES pH 7.5, 500 mM NaCl, 1 mM Benzamidine, 1 mM DTT, 1 mM EDTA) supplemented with 1 cOmplete protease inhibitor tablet/100ml and 0.1 mg/mL lysozyme, then clarified by centrifugation at 40,000 x *g*. Lysates were incubated with 3 mL Glutathione Sepharose 4B for 3 hours, washed and cleaved from immobilized GST fusion tag by overnight incubation with PreScission protease at 4 °C. Finally, the proteins were subjected to size-exclusion chromatography with a HiLoad 16/600 Superdex 75 column equilibrated in 25 mM HEPES pH 7.5 and 150 mM NaCl. Template DNA for GluN2B (vector pCI-EGFP-NR2b) was provided by Andres Barria & Robert Malinow (RRID:Addgene_45447)(Barria & Malinow, 2002), and α-actinin-2 coding sequence by Kristina Djinović-Carugo (Almeida Ribeiro Jr. et al., 2014).

N-terminally 6His-tagged mouse CaMKIIα was cloned into pcDNA3.1 using primers EcoRI_6HisCaMKIIα & CaMKIIα_Term_XhoI for expression in adherent HEK293T cells. Cells were transfected at 60-70% confluency with 10 μg DNA and 60 μg polyethlyenimine (MW 25000) applied to each 10-cm dish (Curtis et al., 2022). The media was replaced with DMEM supplemented with 10% FBS the morning after transfection, and cells were harvested 3 days after transfection. Cells were lysed by Dounce homogenisation in lysis buffer (20 mM HEPES pH 8.0, 20 mM NaCl, 2 mM DTT, 1 mM EDTA, 1 cOmplete protease inhibitor tablet/100 mL) and clarified by centrifugation for 30 min at 45,000 x *g*. CaMKII was initially enriched using anion exchange with Q Sepharose Fast Flow resin (GE healthcare), eluted in high salt buffer (20 mM HEPES pH 7.5, 500 mM NaCl, 2 mM DTT and 1 mM EDTA) following 2 hour incubation. This eluate was exchanged into Ni-NTA buffer A (500 mM NaCl, 25 mM Tris pH 8, 1 mM Benzamidine, 20 mM imidazole) using a HiPrep desalting column (Cytiva) for binding to Ni-NTA agarose. His-CaMKIIα was eluted from the Ni-NTA agarose using a gradient into Ni-NTA buffer B (500 mM NaCl, 25 mM Tris pH 7.5, 1 mM Benzamidine, 300 mM imidazole), and finally exchanged into storage buffer (25 mM HEPES pH 7.5, 150 mM NaCl, 10% w/v glycerol). For expression of CaMKIIα 1-315, the corresponding coding sequence was cloned into pNH-TrxT (gift of Opher Gileadi, RRID:Addgene_26106)(Savitsky et al., 2010) for expression with a Tev-cleavable N-terminal 6His-Trx tag in bacteria using primers pNH-Trx CaMKII_1to315_F & _R. The expression vector was transformed into Rosetta (DE3) pLysS *E. coli*, which were grown in LB and induced with 0.2 mM IPTG at OD_600 nm_ ~ 0.5. Cells were harvested following overnight incubation at 18 °C, then lysed in Ni-NTA Buffer A (25 mM Tris pH 8, 500 mM NaCl, 1 mM benzamidine, 20 mM imidazole) supplemented with 0.1 mg/mL lysozyme and 1 Complete EDTA-free protease inhibitor tablet/100 mL. 6His-Trx-CaMKIIα (1-315) was initially purified by affinity to Ni-NTA agarose, eluting with a gradient into Nickel Buffer B. The eluate was exchanged into anion exchange buffer A (20 mM Tris pH 8.8, 20 mM NaCl, 1 mM DTT, 1 mM EDTA) using a HiPrep 26/10 desalting column to enable binding to a Q Fast Flow column (Cytiva). Protein was eluted using a gradient into anion exchange buffer B (20 mM Tris, pH 8, 500 mM NaCl, 1 mM DTT, 1 mM EDTA). Finally, size exclusion was performed using a HiLoad Superdex 75 column equilibrated in gel filtration buffer (20 mM HEPES pH 7.5, 150 mM NaCl, 1 mM DTT).

Human CaM was expressed and purified as described previously (Patel et al., 2017). Briefly, untagged CaM was expressed using pET28-a in *E. coli* BL21 (DE3) cells grown in AIM. CaM was initially purified by affinity to phenyl sepharose (GE Life Sciences) in the presence of 5 mM CaCl_2_. CaM was eluted in buffer containing 1 mM EDTA then further purified using anion exchange with a Resource Q column (GE Life Sciences) before dialysis into water and lyophilization in a vacuum concentrator.

#### Crystallography

For crystallization of α-actinin-2 EF3-4 with CaMKIIα regulatory segment, peptide corresponding to CaMKIIα positions 294-315 (sequence NARRKLKGAILTTMLATRNFSG, acetylated at N-terminus, amidated at C-terminus) was synthesised by Biomatik at > 95 % purity. α-actinin-2 EF3-4 (16 mg/mL) was mixed with a 2.5-fold molar excess of CaMKIIα peptide in a precipitant solution containing 0.1 M BIS-TRIS pH 6.5, 25% w/v Polyethylene glycol 3,350, and crystals were grown by sitting drop vapor diffusion at 24 °C. Diffraction data was collected to high resolution at Diamond Light Source beamline I04 with x-rays at a wavelength of 0.9795 Å, and at beam line P13 at the PETRA III storage ring (DESY, Hamburg, Germany) with x-rays tuned to 2.0664 Å to amplify the anomalous signal of sulfur. Diffraction data was reduced using DIALS (Winter et al., 2018) and scaled with Aimless (Evans & Murshudov, 2013) before experimental phasing using single-wavelength anomalous diffraction of native sulfur atoms in CRANK2 (Skubak et al., 2018). Refinement was completed in PHENIX (Liebschner et al., 2019). The following residues were omitted from the final model since they could not be clearly resolved in the electron density: EF3-4 892-894 (chain B); CaMKIIα 314-315 (chain C); and CaMKIIα 311-315 (chain D). Full data collection and refinement statistics are provided in **Supplementary Table 1**.

#### Isothermal titration calorimetry

ITC experiments were performed using a MicroCal PEAQ™ (Malvern Panalyticial). To investigate binding between CaMKII peptides and EF1-4/EF3-4, experiments were performed at 25 °C in 25 mM HEPES pH 7.5 and 150 mM NaCl, where 500 μM CaMKII peptides were injected into a cell containing 50 μM EF1-4/EF3-4 using 2 μL injections at 2 min intervals and a constant mixing speed of 750 rpm. Measurements between Trx-CaMKIIα 1-315 and EF3-4 were performed using 30 μM Trx-CaMKIIα 1-315 and 300 μM of EF hands 3-4 and additionally supplementing the buffer with 2 mM MgCl_2_ and 1 mM ADP. To investigate CaMKII-CaM interactions, ITC was performed at 10 °C using 30 μM Trx-CaMKIIα 1-315 - 300 μM CaM (buffer supplemented with 2 mM MgCl_2_, 1 mM ADP, and 1 mM CaCl_2_); and 30 μM CaM - 300 μM CaMKII regulatory peptide (buffer supplemented with 1 mM CaCl_2_). All ITC measurements were collected, baselined and integrated before non-linear least-squares fitting to single binding models using MicroCal origin software.

#### Culture and transfection of primary hippocampal neurons

Primary hippocampal cultures were prepared from E18 Sprague-Dawley rats and plated onto 13 mm coverslips, pre-treated with Poly-L-Lysine (1 mg/mL) at a density of 1×10^5^ cells per coverslip. Neurons were cultured in neurobasal medium supplemented with B27, GlutaMAX and Penicillin/Streptomycin (ThermoFisher Scientific). On DIV10, neurons were transfected with 0.8 μg DNA and 2 μL lipofectamine 2000 (Thermo Fisher) per 13 mm coverslip in 6-well plates using pIRES2-GFP vectors containing N-terminally FLAG-tagged coding sequences for FLAG-α-actinin-2. The vectors were constructed using primers listed in **Supplementary Table 2**. The L854R mutation was introduced into pcDNA3.1-FLAG-α-actinin-2 (747-894) by site-directed mutagenesis with primers L854R_F & _R. After transferring transfected neurons into fresh media on DIV11, *in vitro* culture was continued until chemical LTP/fixing on DIV14.

#### Chemical LTP

Chemical LTP (cLTP) was induced in cultured hippocampal neurons by activating NMDARs with glycine as described previously (Fortin et al., 2010; McLeod et al., 2018). Primary hippocampal neurons were first transferred into control solution (5 mM HEPES pH 7.4, 125 mM NaCl, 2.5 mM KCl, 1 mM MgCl_2_, 2 mM CaCl_2_, 33 mM D-glucose, 20 μM D-APV, 3 μM strychnine, 20 μM bicuculline, 0.5 μM TTX) for 20 min at room temperature (RT). cLTP was induced by incubating for 10 min at RT in cLTP solution (5 mM HEPES pH 7.4, 125 mM NaCl, 2.5 mM KCl, 2 mM CaCl_2_, 33 mM D-glucose, 3 μM strychnine, 20 μM bicuculline, 0.2 mM glycine). Following cLTP induction, neurons were returned to control solution for between 1 and 4 hours before fixation in fixing solution (PBS supplemented with 4 % paraformaldehyde PFA, 4 % sucrose, 0.2% Glutaraldehyde). For PLA assays, neurons were fixed two hours after cLTP induction.

#### Proximity ligation assays and confocal imaging

PLAs were performed using reagents from a Duolink In Situ PLA kit. Fixed neurons were permeabilised for 5 min at RT in PBS supplemented with 1 % BSA/0.1% Triton X-100, and blocked for 1 hour in PBS supplemented with 10 % BSA before overnight incubation at 4°C with primary antibodies (mouse anti-CaMKIIα, 1 in 400 dilution; goat anti-FLAG, 1 in 200 dilution; rabbit anti-GFP, 1 in 300 dilution) in PBS supplemented with 1% BSA. On the following morning, neurons were incubated with the corresponding Duolink probes (anti-goat PLUS and anti-mouse MINUS) and goat anti-rabbit Alexa Fluor-405 (ThermoFisher) at 37 °C for 1 hour. Probes were ligated at 37°C for 30 min and signals were amplified at 37 °C for 100 min. All neurons were imaged using ZEN software with a Zeiss LSM 700 inverted microscope using a 60x oil objective (NA=1.40). For PLA experiments, images were collected using 405 nm excitation/421 nm emission for GFP, and 594 nm excitation/619 nm emission for PLA puncta. For imaging neurons transfected with α-actinin-2 EF1-4 disruptors, intrinsic GFP fluorescence was imaged using 488 nm excitation/509 nm emission. Data were obtained from at least three independent neuronal cultures unless otherwise stated. Images were analysed using NeuronStudio software (Icahn School of Medicine at Mount Sinai) to determine spine width and morphology; and the Distance Analysis (DiAna) plugin (Gilles et al., 2017) for ImageJ (NIH) to identify PLA puncta.

#### Magnetic bead pull-down assays

For each CaMKIIα pull-down assay, 0.25 μg of Pierce Glutathione Magnetic Agarose Beads were charged with 4 μg GST or GST-EF1-4 in basic binding buffer (25 mM HEPES pH 7.5, 150 mM NaCl, 0.05 % Tween-20, 10 mM MgCl_2_, 0.5 mM ADP, 2 mM DTT) for 2 hours at 4 °C. Protein mixtures containing CaMKII, CaM, and GluN2Bc (as appropriate) were separately pre-incubated for one hour in basic binding buffer supplemented with CaCl_2_. These mixtures were diluted with equal volumes of basic binding buffer containing 4 % BSA to achieve final concentrations of 2 mM CaCl_2_, 2 % BSA, 0.1 μM CaMKIIα, 3 μM CaM, and 1.5 μM GluN2Bc as appropriate. For each assay, 100 μL protein mixture was incubated with GST/GST-EF1-4 magnetic beads for 20 mins before the reactions were supplemented with either 10, 2.5, or 1.8 mM EGTA. Free Ca^2+^ concentrations were estimated using maxchelator (Bers et al., 2010). Following 20 min further incubation, each pull-down was washed with 3 x 500 μL basic binding buffer supplemented with 2 mM CaCl_2_ and the appropriate concentration of EGTA. Proteins were eluted from the beads by incubation with 50 μL 1×LDS loading buffer (5 min heating at 65 °C). CaMKII and GST/GST-EF1-4 were detected using immunoblotting with mouse anti-CaMKIIα and rabbit anti-GST antibodies.

#### Statistical analysis

Data were assessed for normality using Kolmogorov-Smirnov testing. Normally distributed data were analyzed using unpaired two-tailed Student’s t-tests whereas Mann-Whitney tests were applied for non-parametric data. \**p* < 0.05; \**\*p* < 0.01; \**\*\*p* < 0.001

## Data availability

Coordinates and structure factors have been deposited with the RCSB Protein Databank for the EF3-4 – CaMKIIα regulatory segment peptide complex with accession ID 6TS3.

## Author contributions

A.C. imaged neurons, carried out isothermal titration calorimetry, developed expression vectors, performed protein purification, analyzed data, and prepared figures. J.Z. solved the crystal structure, developed expression vectors, and performed protein purification. C.P. developed expression vectors. M.G.G. conceived the study, performed magnetic bead pull-down experiments, analyzed data, and wrote the manuscript with input from all of the authors.

## Acknowledgements

MGG is a Wellcome Trust and Royal Society Sir Henry Dale fellow (104194/Z/14/Z). This work was supported by BBSRC project grant BB/N015274/1, and a BBSRC studentship to AC. We are grateful to Opher Gileadi, Andres Barria, Robert Malinow, and Kristina Djinović-Carugo for providing DNA vectors in support of this work.

## Competing Interests

There are no financial or non-financial competing interests

## Ethics

Experiments involving rats were performed in accordance with the United Kingdom Animals Act, 1986 and within University College London Animal Research guidelines overseen by the UCL Animal Welfare and Ethical Review Body under project code 14058.

## Supplementary Information

**Figure 1-figure supplement 1.**
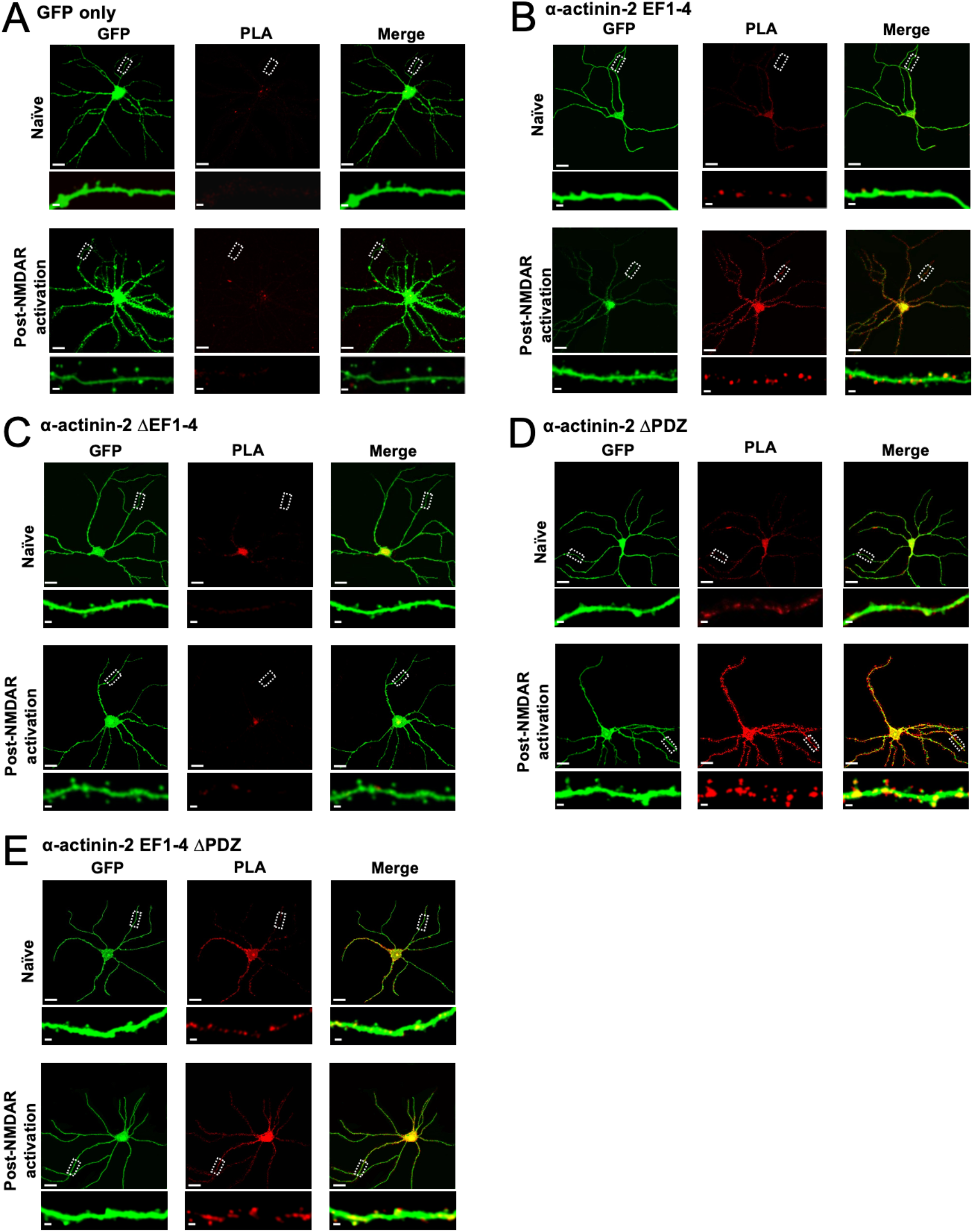
PLA imaging of CaMKIIα association with fragments of α-actinin-2. Each panel shows anti-GFP immunofluorescence (left columns) and anti-CaMKIIα/anti-FLAG PLA puncta (middle columns) in primary hippocampal neurons either before (upper row) or after (lower row) NMDAR activation. The images correspond to transfections with GFP alone (A), or GFP in combination with either α-actinin-2 EF1-4 (B), ΔEF1-4 (C), ΔPDZ (D), or EF1-4 ΔPDZ (E). Scale bars correspond to 20 *μm* (whole neuron images) and 1 *μm* (dendrite close-ups).

**Figure 2-figure supplement 1.**
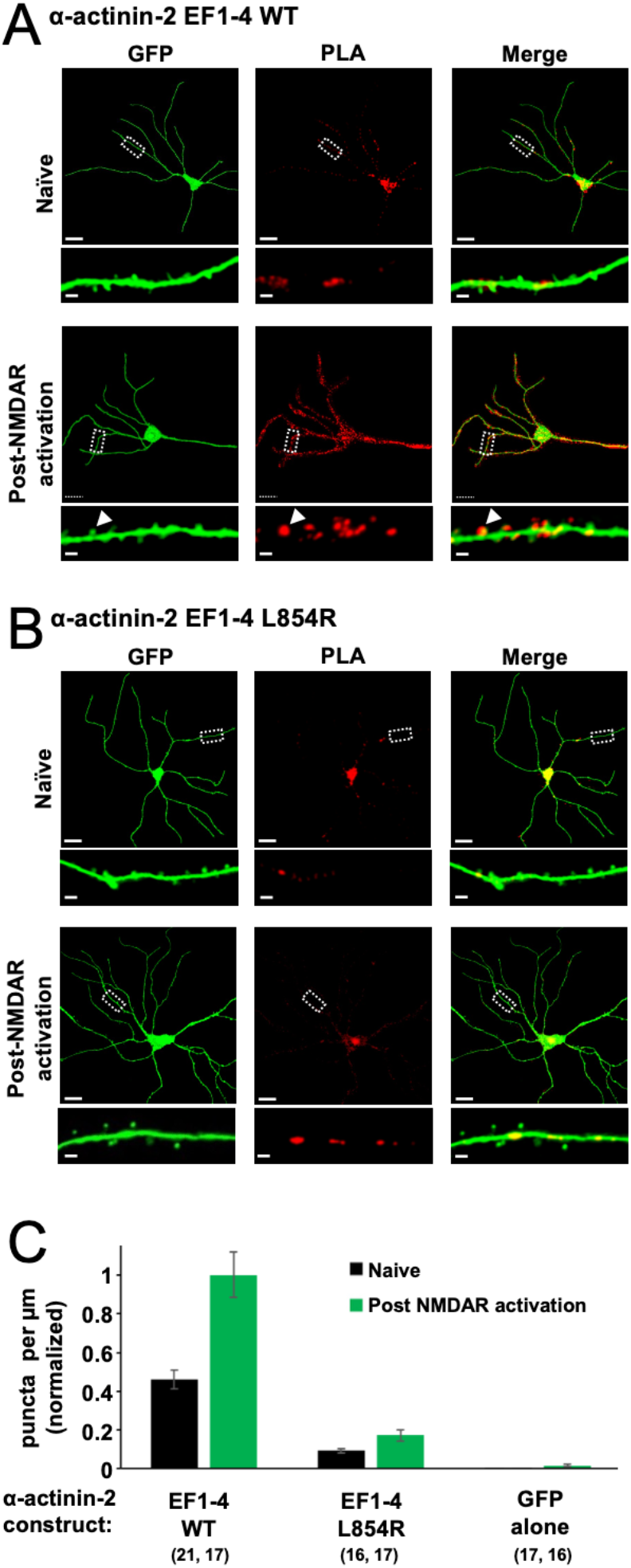
PLA imaging of CaMKIIα association with α-actinin-2 EF1-4 disruptors. The first two panels show imaging following transfections with GFP in combination with either WT (A) or L854R (B) α-actinin-2 EF1-4. Each panel shows anti-GFP immunofluorescence (left columns) and anti-CaMKIIα/anti-FLAG PLA puncta (middle columns) in primary hippocampal neurons either before (upper row) or after (lower row) NMDAR activation. Scale bars correspond to 20 *μm* (whole neuron images) and 1 *μm* (dendrite close-ups). (C) Quantitation of dendritic PLA puncta density before (black) and after (green) NMDAR activation. Data are normalised to EF1-4 WT post NMDAR activation. The number of neurons analysed for each construct is shown in parentheses. PLA puncta density was reduced by 80 % pre-activation (*p*=2.4×10^-7^) and 83 % post NMDAR-activation (*p*=2.0×10^-7^) in neurons expressing EF1-4 L854R compared to the WT variant.

**Figure 2-figure supplement 2.**
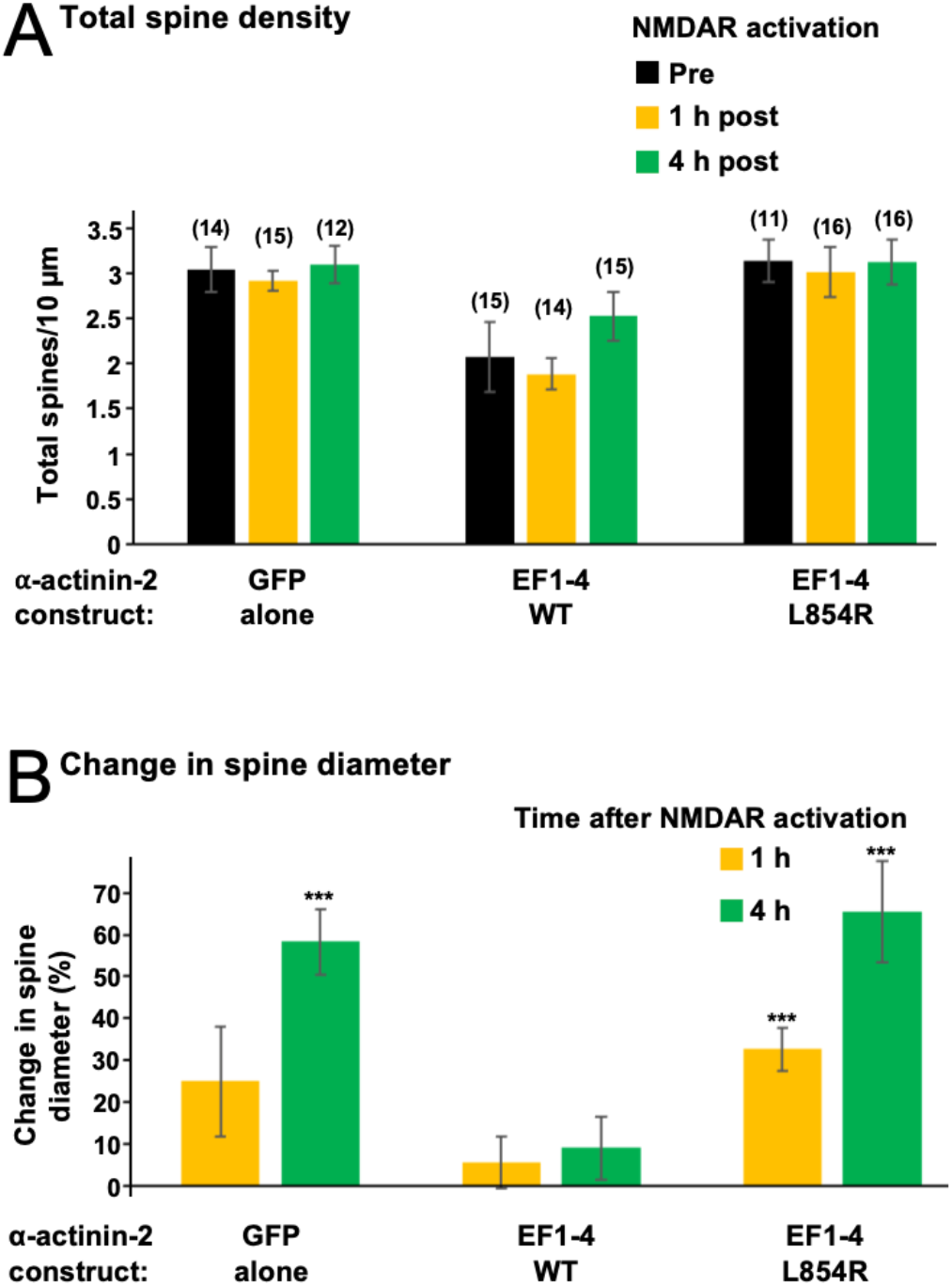
Effect of EF hand disruptors on total spine density and average spine diameter. (A) Total spine density in primary hippocampal neurons transfected with either GFP alone, or GFP in combination with either WT or L854R α-actinin-2 EF1-4. Mean spine number ± SE per 10 μm dendrite is shown before NMDAR activation (black), and 1 hour (orange) and 4 hours (green) after activation. Numbers of neurons analysed for each condition are shown in parentheses. Before NMDAR activation, neurons expressing EF1-4 WT had 32 % fewer spines than those expressing GFP alone (p=0.0013); and 34 % fewer total spines than those expressing the L854R variant (p=5.33×10^-5^). (B) Changes in spine diameter (mean ± SE) are shown either 1 (orange) or 4 (green) hours after NMDAR activation.

**Figure 3-figure supplement 1.**
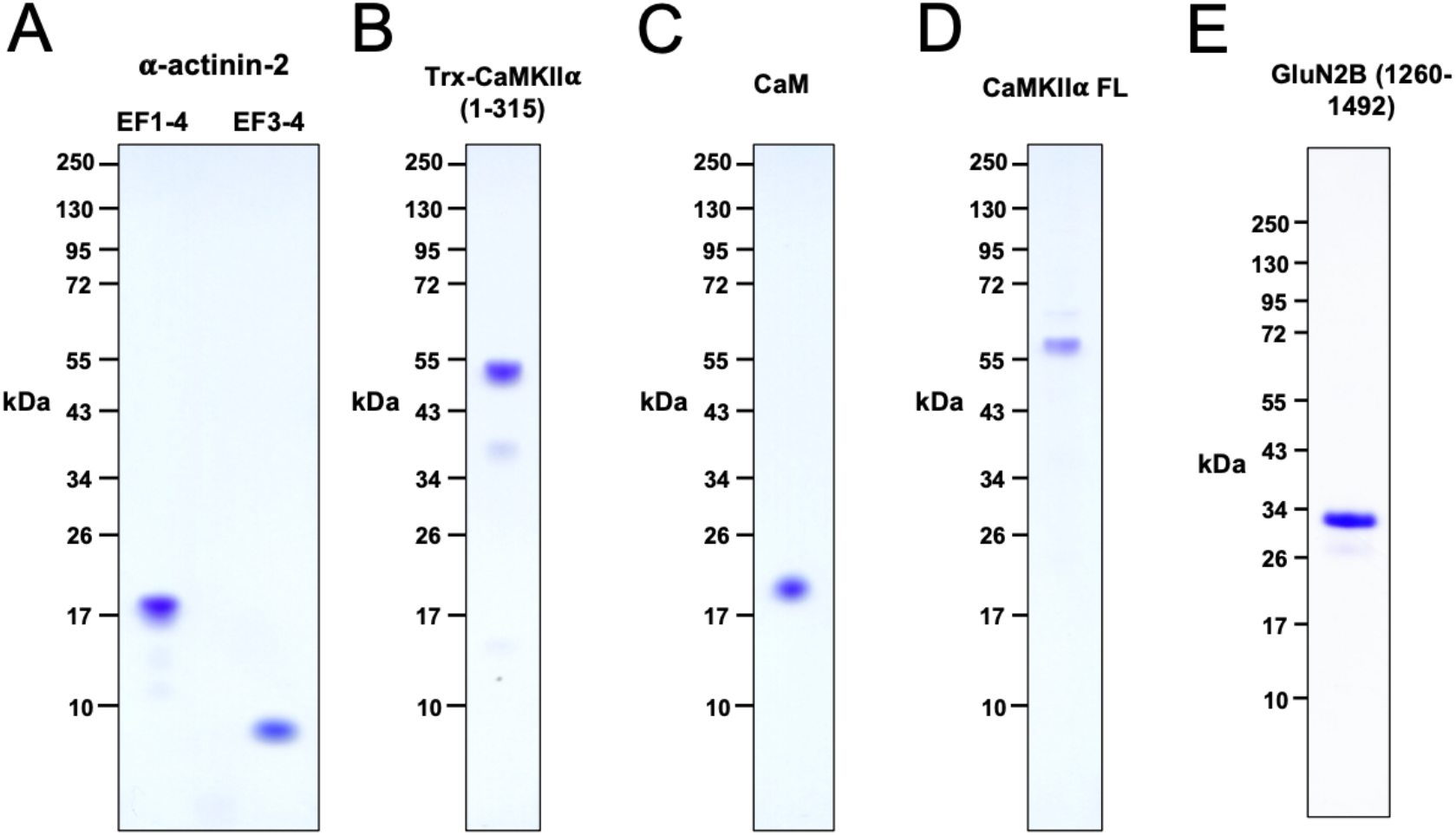
Purified proteins. Coomassie-stained 4-12 % polyacrylamide gels are shown for purified EF1-4/EF3-4 fragments of α-actinin-2 (A), Trx-CaMKIIα 1-315 (B), CaM (C), full-length CaMKIIα (D), and GluN2B 1260-1492 (E).

**Figure 4-figure supplement 1.**
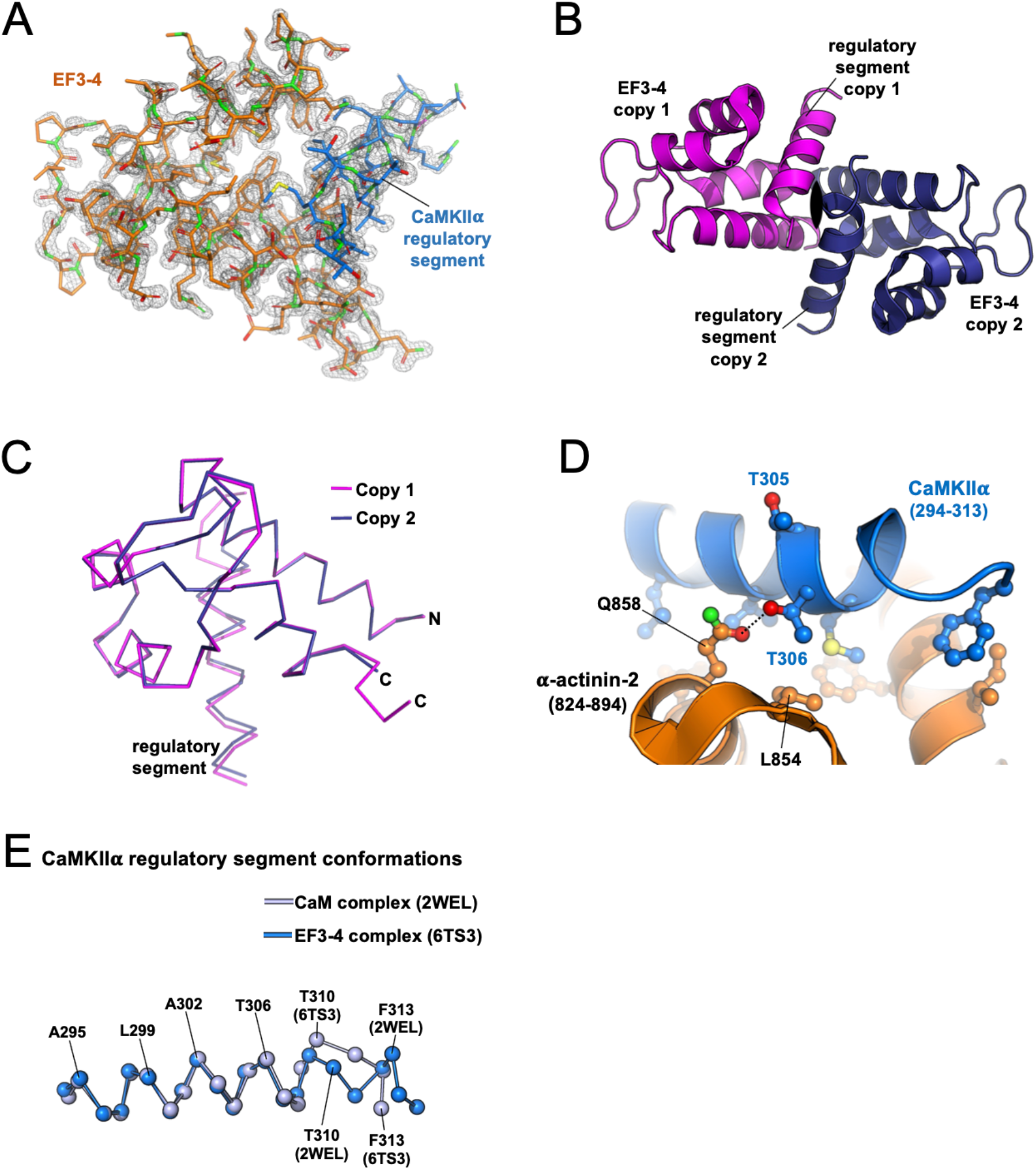
Additional features of the α-actinin-2 EF3-4 - CaMKIIα regulatory segment crystal structure. (A) 2F_o_-F_c_ electron density map is contoured at 1.5 *σ* and clipped within 1.5 Å of the polypeptides (chain A & C). (B) Asymmetric unit showing two copies of the EF3-4 – CaMKIIα regulatory segment complex related to each other by a two-fold rotational symmetry axis. (C) Superimposition of the two copies of the complex. The first copy (chains A & C) is shown in magenta; the second copy (chains B & D) in dark blue. The two copies align with RMSD = 0.202 Å for all atoms encompassing α-actinin-2 822-891 & CaMKIIα 294-310. (D) Close-up highlighting the positions of CaMKIIα threonines 305 and 306 in the complex.

**Figure 4-figure supplement 2.**
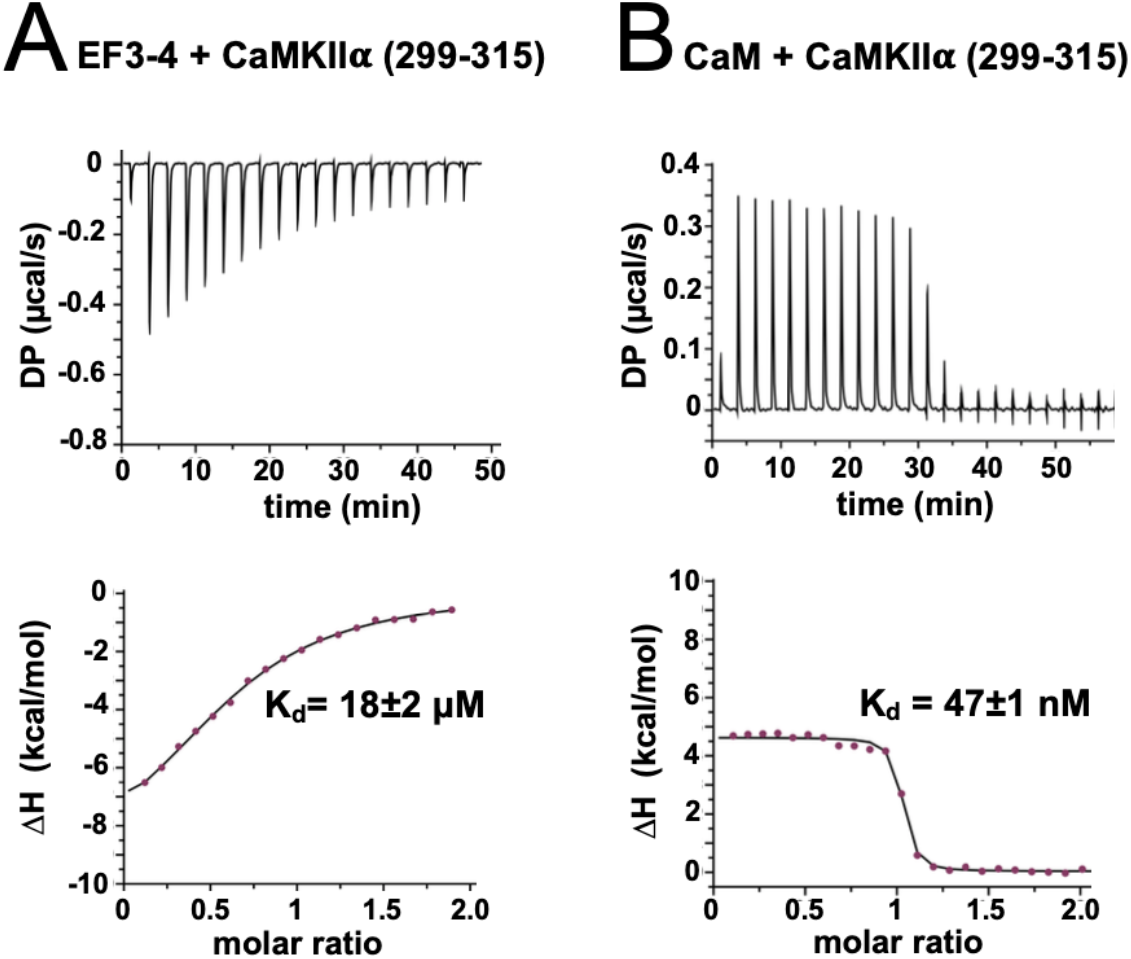
Calorimetry with CaMKIIα 299-315 peptide. Representative isotherms are shown for titrations of the peptide with (A) α-actinin-2 EF3-4 and (B) CaM. The top sub-panels show the raw power output (μcal/s) per unit time; the bottom sub-panels show the integrated data including a line of best fit to a single site binding model. Stated K_d_ values are averages from three replicates.

**Figure 5-figure supplement 1.**
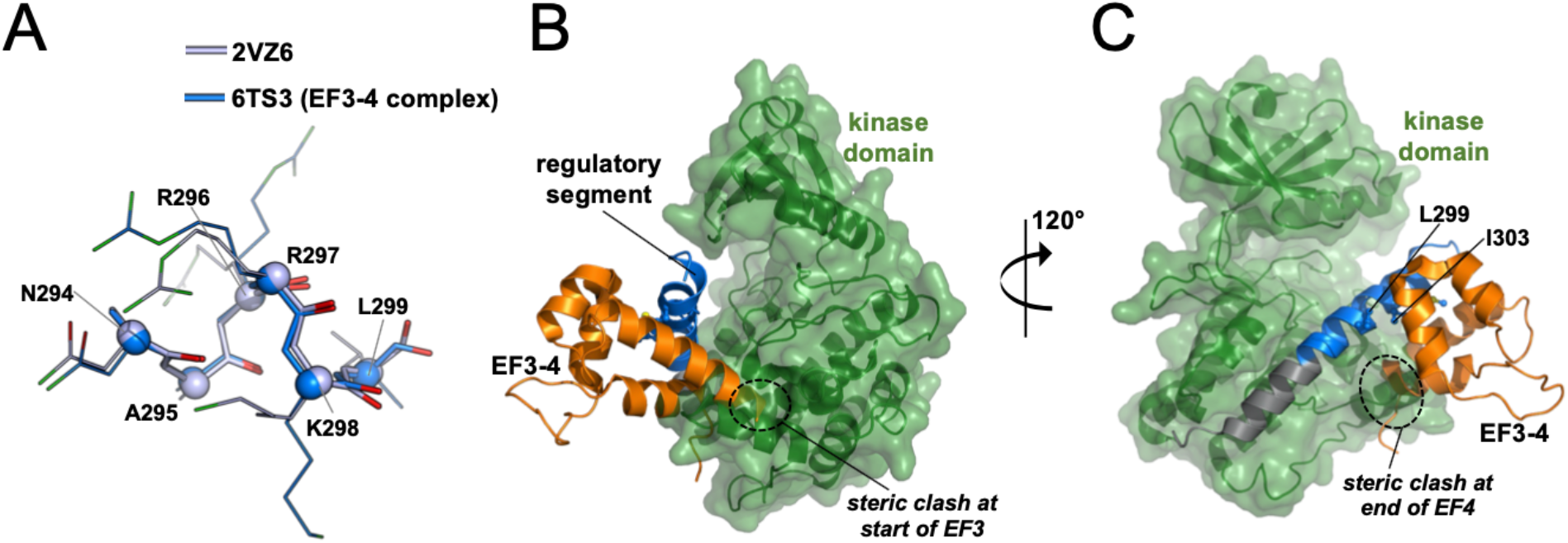
Modelling relative orientations of kinase and EF3-4 domains relative to the regulatory segment. (A) Alignment of all atoms for CaMKIIα positions 294-299 in crystal structures 2VZ6 and 6TS3. Cα atoms are shown as spheres. Panels (B) and (C) show two views of the superimposition of the α-actinin-2 – regulatory segment complex (6TS3) onto the structure of CaMKIIα (3SOA – *β*7 linker and hub domain are not shown). The structures were aligned through positions 294-305 of the regulatory segment.

**Supplementary Table 1.** Data Collection & Refinement Statistics.

| | CaMKII peptide +<br>α-actinin-2 EF3-4 | CaMKII peptide + α-actinin-2<br>EF3-4 (native sulfur<br>anomalous scattering) |
| --- | --- | --- |
| PDB Code | 6TS3 |  |
| Data Collection |  |  |
| X-ray wavelength<br>(Å) | 0.9795 | 2.0664 |
| Resolution range<br>(Å) | 44.46-1.28 | 44.52-2.38 |
| Space group | P1211 | P1211 |
| Cell dimensions |  |  |
| a, b, c (Å) | 43.35, 47.27, 47.47 | 43.35, 47.27, 47.47 |
| $\alpha$, $\beta$, $\gamma$ (°) | 90.0, 110.53, 90.0 | 90.0, 110.53, 90.0 |
| Unique Reflections | 46303 (2152) | 7653 (521) |
| Multiplicity | 6.6 (5.2) | 18.1 (11.8) |
| Completeness (%) | 99.8 (95.4) | 94.5 (66.8) |
| Mean I/$\sigma$(I) | 19.7 (2.4) | 52.2 (26.8) |
| R<sub>pim</sub> | 0.015 (0.208) | 0.011 (0.019) |
| CC<sub>1/2</sub> | 0.999 (0.94) | 0.999 (0.999) |
| Anomalous<br>completeness (%) |  | 92.1 (57.4) |
| Anomalous<br>multiplicity |  | 9 (6.4) |
| Copies per ASU | 2 | 2 |
| Refinement statistics |  |  |
| R<sub>work</sub> | 0.183 |  |
| R<sub>free</sub> | 0.199 |  |
| R.m.s. deviations<br>of bond lengths (Å) | 0.005 |  |
| R.m.s. deviations<br>of bond angles (Å) | 0.698 |  |
| Ramachandran |  |  |
| Favored (%) | 99.4 |  |
| Allowed (%) | 0.6 |  |
| Outliers (%) | 0 |  |

**Supplementary Table 2.**
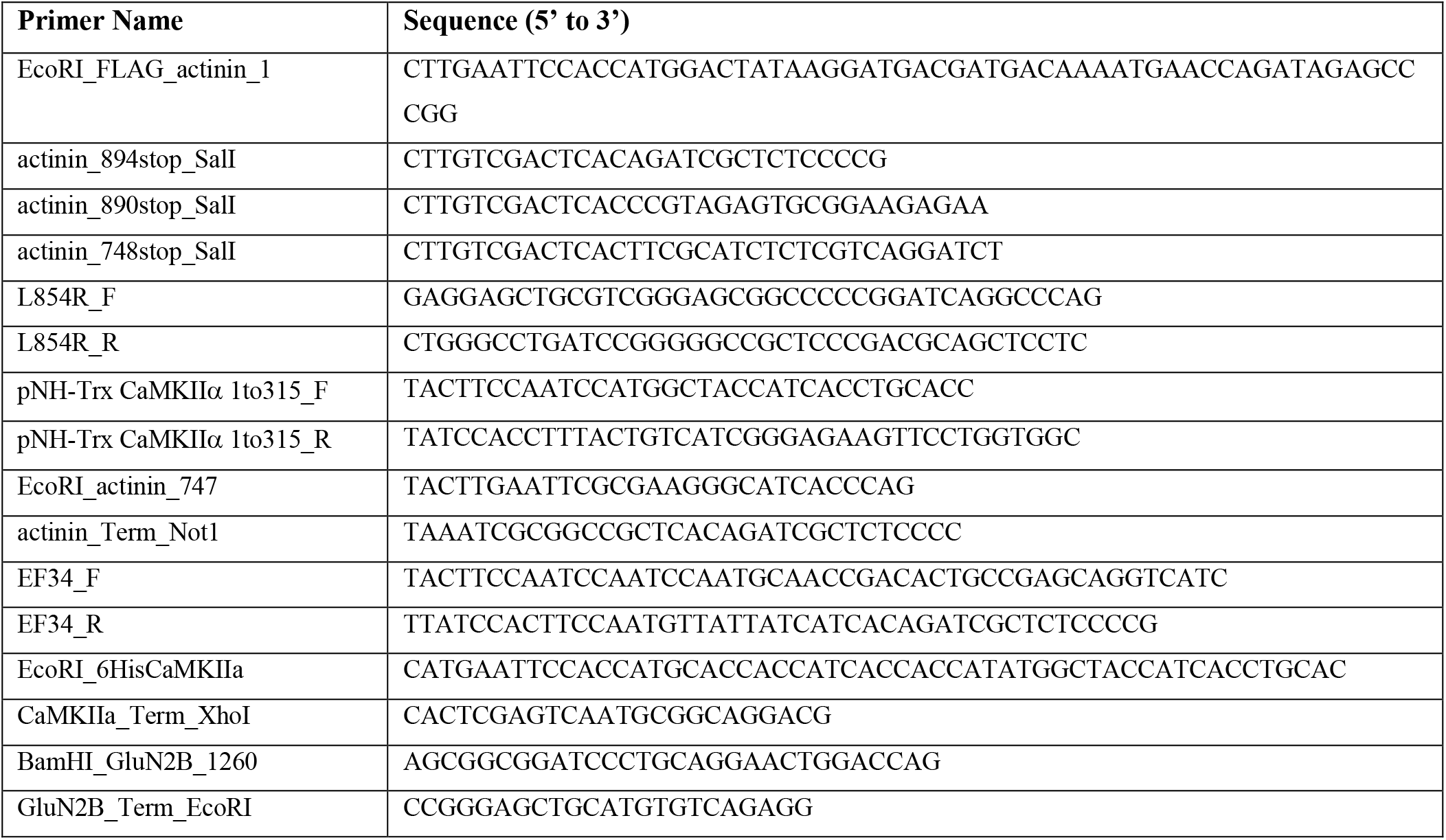
Oligonucleotide primers sequences.

## Legends for source data linked to figures

**Figure 1-source data 1**. Full PLA dataset.

**Figure 2-source data 1**. Full spine classification dataset.

**Figure 2-figure supplement 1-source data 1**. Full PLA data for disruptor experiments.

**Figure 3-source data 1**. Full ITC dataset.

**Figure 3-figure supplement 1-source data 1**. Uncropped Coomassie-stained gels.

**Figure 3-figure supplement 1-source data 2**. Raw image for panels A-D.

**Figure 3-figure supplement 1-source data 3**. Raw image for panel E.

**Figure 5-source data 1**. Densitometry breakdown for CaMKII pull-down experiments.

**Figure 5-source data 2**. Uncropped immunoblots.

**Figure 5-source data 3**. Raw image for anti-CaMKII immunoblot.

**Figure 5-source data 4**. Raw image for anti-GST immunoblot.

## Notes

### Competing Interest Statement

The authors have declared no competing interest.

